# An integrase toolbox to record gene-expression during plant development

**DOI:** 10.1101/2022.09.16.508262

**Authors:** Sarah Guiziou, Cassandra J. Maranas, Jonah C. Chu, Jennifer L. Nemhauser

**Affiliations:** Department of Biology, University of Washington, Seattle, Washington 98195, USA

**Author notes:** Author for communication.

## Abstract

There are many open questions about the mechanisms that coordinate the dynamic, multicellular behaviors required for organogenesis. Synthetic circuits that can record *in vivo* signaling networks have been critical in elucidating animal development. Here, we report on the transfer of this technology to plants using orthogonal serine integrases to mediate site-specific and irreversible DNA recombination visualized by switching between fluorescent reporters. When combined with promoters expressed during lateral root initiation, integrases amplified reporter signal and permanently marked all descendants. In addition, we have developed a suite of methods to tune the threshold for integrase switching, including: RNA/protein degradation tags, a nuclear localization signal, and a split-intein system. These tools improved the robustness of integrase-mediated switching with different promoters and the stability of switching behavior over multiple generations. This integrase toolbox can be used to build history-dependent circuits to decode the order of expression during organogenesis in many contexts.

## Introduction

Biologists have long been fascinated by the molecular pathways that support the development of complex multicellular organisms. Plants are particularly intriguing subjects to study, as the development programs that start in their embryos persist throughout their lifespan, strongly influenced by environmental cues. The growing environmental pressures resulting from climate change make this adaptability increasingly important^1^. Better understanding of the mechanisms that underlie plant developmental plasticity will help guide the engineering of traits that can face current and future challenges^2^.

To fully understand the molecular trajectory underlying fate transitions that enable *de novo* organogenesis and regeneration in plants, we need methods that can sense and relay information in a manner that can be dynamically and quantitatively read out by an observer. Current methods enable precise quantification of DNA^3^, RNA^4^, and proteins^5^ allowing the capture of a snapshot of the molecular state of studied organisms. Combining these approaches with single-cell methods has led to the discovery of new plant cell types and a more detailed view of cell-fate transitions^6–12^.

A challenge of current single-cell methods is that they require the destruction of samples, and, therefore, do not allow for real-time readouts, reports from the same sample across multiple timepoints, or preserve spatial relationships. With recent advances in high-throughput and high-precision microscopy, fluorescent reporters and sensors have allowed imaging at cellular resolution of transcription level, protein and molecule concentration and localization in a continuous manner in their native context^13^. However, detection is limited to a reduced amount of information at a time due to the limited number of fluorescent tags and to short timescales due to photobleaching and stress to the organisms. Recently, the development of synthetic, DNA-based recording systems have overcome some of the technical challenges of ‘omic and microscopy techniques, allowing the sensing and relaying of multiple signals simultaneously during animal development (reviewed in ^2^).

Serine integrases, used by bacteriophages to mediate their own integration into the bacterial genome, were critical to the success of one of the most promising synthetic recorders^14^. In a synthetic system, serine integrases are used to invert or excise DNA in a site-specific and irreversible manner, referred to here on as an integrase switch. The integrase recognizes two DNA sites of around 40bp known as attB and attP sites. If the sites are in the same orientation, the DNA region between them is excised and if the sites are in the opposite orientation, the region is inverted. Gene-regulatory parts, such as promoters or terminators, can be placed between integrase sites to mediate a specific gene expression pattern dependent on integrase expressions. Complex genetic circuits have been developed using integrases, implementing Boolean logic (in bacteria^15, 16^, mammalian cells^17^, and plant protoplasts^18^), history-dependent logic (in bacteria^19, 20^), and cell-lineage tracing (in animals^14^). Serine integrases can also be used to induce expression of toxic genes at a specific time, to mediate site-specific DNA integration^21^, and has been also used for multi-part in vitro cloning^22^.

To date, serine integrases have not been used extensively in plant systems, although they have been shown to work in principle in *Arabidopsis*^23^, *Nicotiana benthamiana*^24 21^, barley^25^ and wheat^26^. One study in *N. benthamiana* used a recombination directionality factor (RDF), which when combined with the integrase, allowed reversing of the integrase reaction^24^. In addition, tyrosine integrases were recently used to implement logic circuits in *Arabidopsis*^18^. Here, we developed a toolbox of well-characterized parts to build synthetic circuits in *Arabidopsis* using PhiC31 and Bxb1 integrases. We expressed integrases from well-characterized transcription factors essential for lateral root development as a test-case for building synthetic recorders. To optimize the specificity and robustness of our tools when using different promoters, we built and tested a variety of methods to tune the threshold for integrase switching—tools that could be used to tune the activity of any protein of interest. Finally, we characterized two methods that allow for further fine-tuning of the timing and level of integrase activity: split-intein-integrase and estradiol-inducible integrase. Collectively, these new, modular parts make it possible to record gene expression at specific times and spaces during plant development, as well as contribute to an accelerated design-test-build-learn cycle for other plant synthetic biology devices.

## Results

### Orthogonal and efficient DNA switches in Arabidopsis

Our first goal was to test the efficiency of three serine integrases (PhiC31, Bxb1, and Tp901) in *Arabidopsis* transgenic lines. To do this, we needed two constructs: the target and the integrase. These two constructs cannot be on the same plasmid, as even low levels of integrase produced in bacteria during cloning will enable target site inversion. For our target construct, we used a constitutive promoter (pUBQ10; ^27^) flanked by integrase sites positioned between two reporter genes: the mScarlet and mTurquoise2 (mTurq) fluorescent proteins (Fig. 1a). To reduce the likelihood of bidirectional expression from strong promoters like the Cauliflower Mosaic Virus 35S promoter (p35S), and to more closely match the expression level of most developmentally-relevant genes, we opted to use the promoter of PROTEIN PHOSPHATASE 2A SUBUNIT 3 (pPP2AA3), a gene which has been widely used as a qPCR control due to its constitutive nature and medium expression level^28^. If expressed and functional, the integrase should mediate inversion of the promoter resulting in the switch of expression from mTurq to mScarlet. Targets containing either PhiC31 or Bxb1 integrase sites strongly expressed mTurq and not mScarlet in roots and leaves (Fig. 1b). When either the PhiC31 or Bxb1 integrase was constitutively expressed alongside the target with its cognate integrase sites, we observed exactly the opposite reporter expression (strong mScarlet and no mTurq) in all tissues. Moreover, we confirmed that PhiC31 and Bxb1 integrases are orthogonal to one other, as Bxb1 integrase does not mediate an integrase switch in the PhiC31 target line nor *vice versa* (Fig. 1c). The Tp901 integrase, known to be less efficient than Bxb1 and PhiC31^29^, did not cause any switching in targets carrying its target sequences, even with strong promoters (p35S and pUBQ10), and codon optimization (Fig. S2). We also tested a switch using YFP and Luciferase reporters that had been used previously in *N. benthamiana* ^24^, and confirmed that the pPP2AA3 promoter allows constitutive integrase switch with this target as well (Fig. S1).

**Figure 1:**
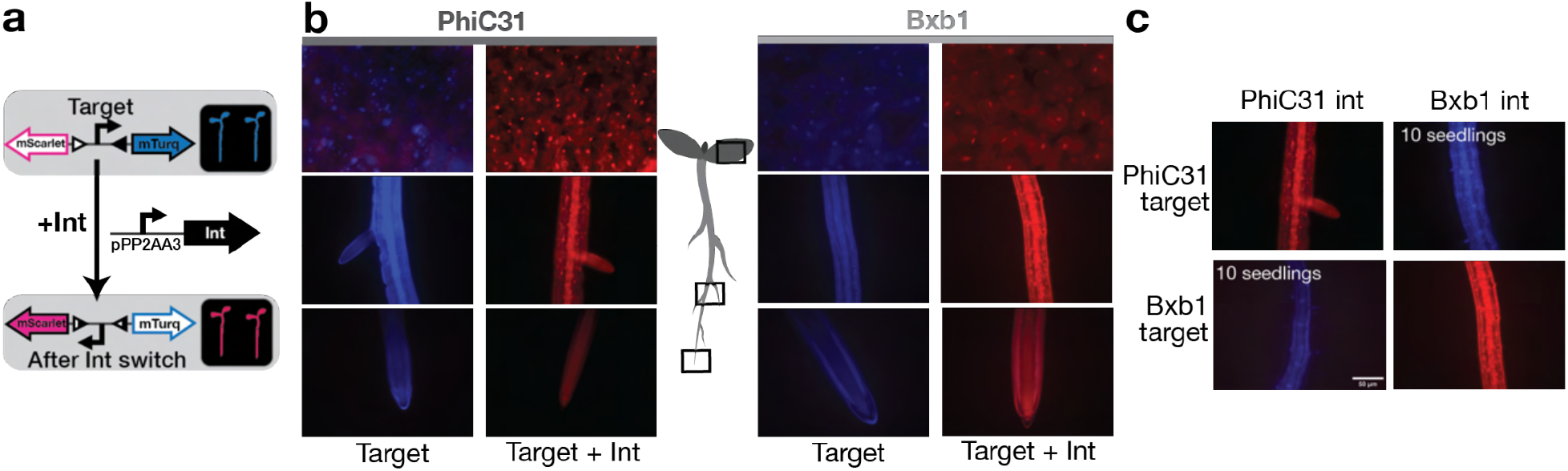
Integrase mediates orthogonal DNA-switch in *Arabidopsis*. (a) Design of the integrase target. The target is composed of two integrase sites (triangles) surrounding a constitutive promoter (pUBQ10) and two fluorescent reporters (mTurquoise2 and mScarlet). In absence of integrase, mTurquoise2 is expressed. In presence of integrase, the integrase mediates inversion of the DNA between the integrase sites, inverting the promoter, and leading to mScarlet expression. The expression of the integrase is mediated by the constitutive promoter pPP2AA3. (b) Constitutive integrase switch characterization. On the left side, *Arabidopsis* seedlings with PhiC31 target alone and PhiC31 target with pPP2AA3::PhiC31 construct. On the right side, Bxb1 target alone, and Bxb1 target with pPP2AA3::Bxb1 construct. Microscopy images are an overlay of mTurq (in blue) and mScarlet (in red) fluorescence, from top to bottom are images of the leaf, a lateral root, and the root tip. (c) Orthogonality test of the integrase switch. Each integrase target line was transformed with both integrases. Microscopy images are overlays of mTurq (in blue) and mScarlet (in red) fluorescence, and are representative images (n=10 seedlings screened).

To record developmental events, switching of the integrase targets must be consistently restricted to a narrow range of time and space. For the next round of design and testing, we focused on the PhiC31 integrase and lateral root development, a well-characterized example of *de novo* organogenesis^30^. Lateral root development is a good model for applying new tools to study gene expression because, it is a well-studied pathway with defined transcriptional control points^31^. The density and placement of lateral roots are also features of plant architecture that are linked to climate resilience and therefore a strong candidate for synthetic engineering^32^. Lateral roots initiate from a small population of founder cells at the xylem pole of the pericycle layer^33^, and follow a fairly stereotyped pattern through morphogenesis^34^.

As test drivers for integrase expression, we selected the promoters of several well-studied transcription factors expressed in the early stages of lateral root initiation: AUXIN RESPONSE FACTOR 7 (ARF7)^35^, AUXIN RESPONSE FACTOR 19 (ARF19)^35^, LATERAL ORGAN BOUNDARIES DOMAIN 16 (LBD16)^36^, and GATA TRANSCRIPTION FACTOR 23 (GATA23)^37^. Because the integrase switch is heritable, we would expect that if the integrase is expressed in lateral root founder cells and works efficiently, all cells in the new root should be in the switched state as well. Simply put, all of the cells in the main root should express mTurq, while all of the cells of the lateral roots should express mScarlet. We characterized approximately 20 independent transformants (T1s) per integrase construct, and categorized each seedling by the following categories: (1) No-switch: expression of mTurq only; (2) LR-only: expression of mScarlet in lateral root only; or (3) Non-exclusive: any expression of mScarlet in the main root. For pARF19 and pGATA23, a majority of the T1s was switched in the lateral root only (81% and 92% of the seedlings, respectively) (Fig. 2a), showing that the integrase switch can record the transcription of a development-related gene. Additionally, this data prove that the integrase system can faithfully trace cell lineage, as all cells, even those in fully emerged lateral roots, continued to express mScarlet only.

**Figure 2:**
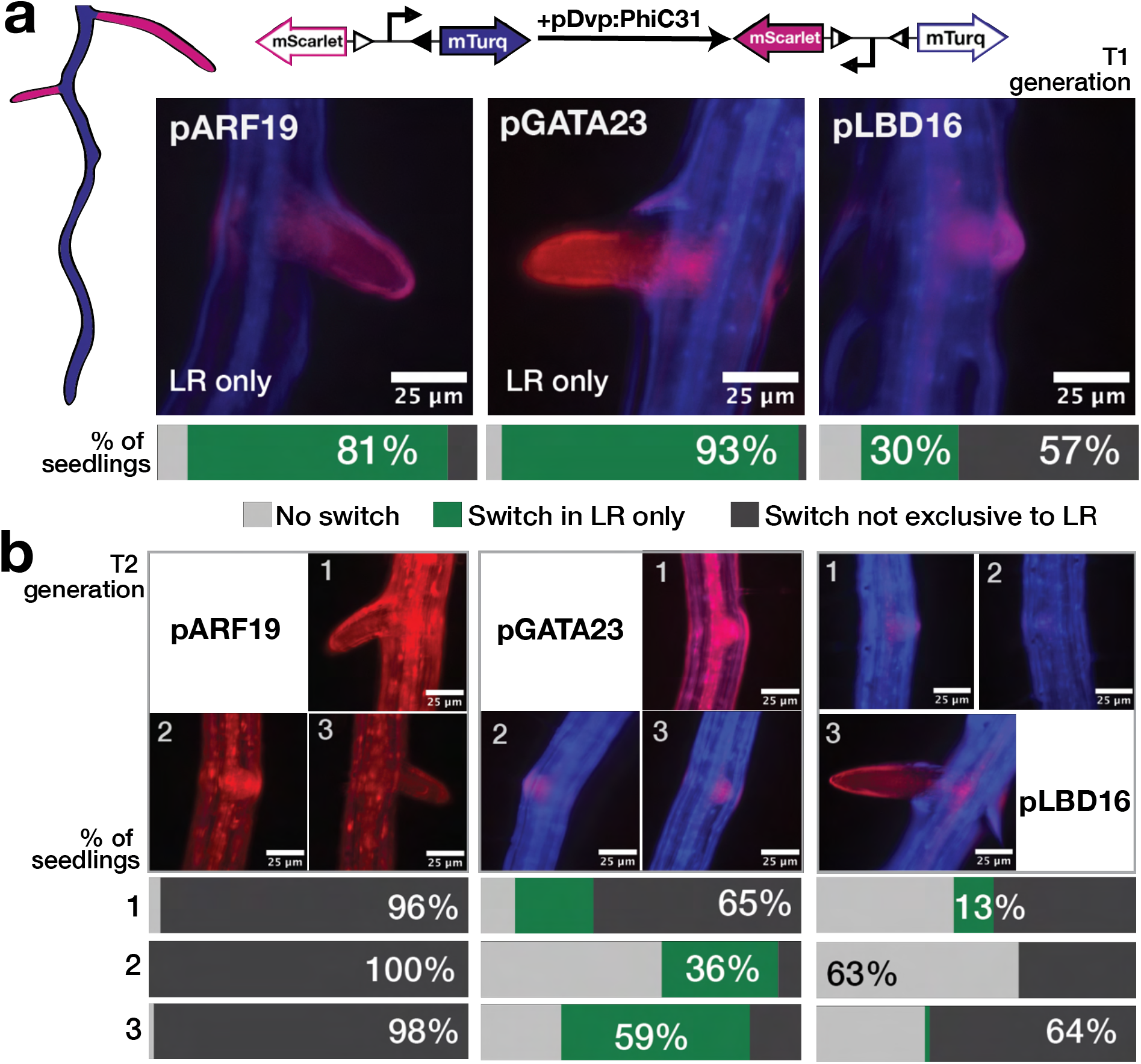
PhiC31 integrase switch under the control of developmental promoters. **(a)** Developmental (Dvp) promoters drove the expression of PhiC31 integrase with a target that switches from mTurquoise2 to mScarlet when the integrase is active. Target lines were transformed with the integrase constructs, and at least 20 T1 seedlings per integrase constructs were characterized. Representative images of emerged lateral roots are shown for each promoter-integrase construct, as well as a bar representing the percentage of seedlings in each phenotypic category: no switch (light gray), switch in LR only (green), or switch not exclusive to LR (dark gray). **(b)** Characterization of T2 seedlings. For each construct, we selected 3 T1 lines with an LR-only switch phenotype, and characterized 20 T2 seedlings per T1 line. For each T1 line, a representative T2 seedling is shown above bar graphs displaying the percentage of seedlings in each phenotypic category. The percentage of the phenotype represented by the T2 image is displayed numerically on the relevant portion of the graph.

Other promoters did not fulfill the specifications. For pLBD16, we observed only 30% LR-only seedlings, while 57% of T1s showed non-exclusive expression of mScarlet (Fig. 2a). Most of the seedlings in the non-exclusive category (70%) did not display switching in the entire seedling, but instead had mScarlet expression in a few cells in the vasculature in addition to the lateral root (Fig. S3). This “weak” non-exclusive switching pattern (Fig. S3) corresponds to the known expression pattern of *LBD16*^38–40^. For *ARF7*, 79% of seedlings were switched in all tissues of the root (Fig. S4). Roughly half of the non-exclusive seedlings showed a full switch and half showed some expression of mTurq (in addition to mScarlet) in the entire root. This result is consistent with *ARF7* being expressed in other tissues and other times of development^41^.

Our next question was whether the integrase system would remain robust over subsequent generations, or whether some low level of leakiness would lead to plants where every cell was in the switched state. For these experiments, we selected three T1 lines where PhiC31 was driven by pARF19, pGATA23 or pLBD16, and which were characterized as having LR-only switching. From each line, we characterized 20 progeny (T2 seedlings). In all cases, we observed a decrease in LR-only seedlings in the T2 generation (Fig. 3b). For pGATA23 T2s, we observed LR-only switches, but at a lower proportion than in T1s, and obtained seedlings displaying no-switch and non-exclusive switches. This pattern is not surprising, as in the T1 generation, plants are hemizygous for the integrase transgene insertion events, meaning that some T2s may end up with no integrase and others may have different numbers of insertions leading to a range of expression levels. For *ARF19*, the majority of T2 seedlings are fully switched (96%, 100%, 98%), also consistent with an increased dosage of integrase in many of the T2s. The lack of no-switch category T2s for this construct suggests that the integrase expression may be happening during gamete development and then transmitted to all of the cells in the T2 generation. For pLBD16 T2s, most of the seedlings are either no-switch or non-exclusive, with 66% or more of the non-exclusive seedlings having a weak switch in the main root similar to the T1 generation (Fig. S3). To obtain a robust, cell-type specific switch, the expression level of the gene of interest in those cells should be significantly greater than that in other cell types. With *LBD16*, it seems that the expression level in LR cells is similar enough to that in the phloem pole pericycle (BAR Webservices, ^38^) to make LR-only switching rare.

**Figure 3:**
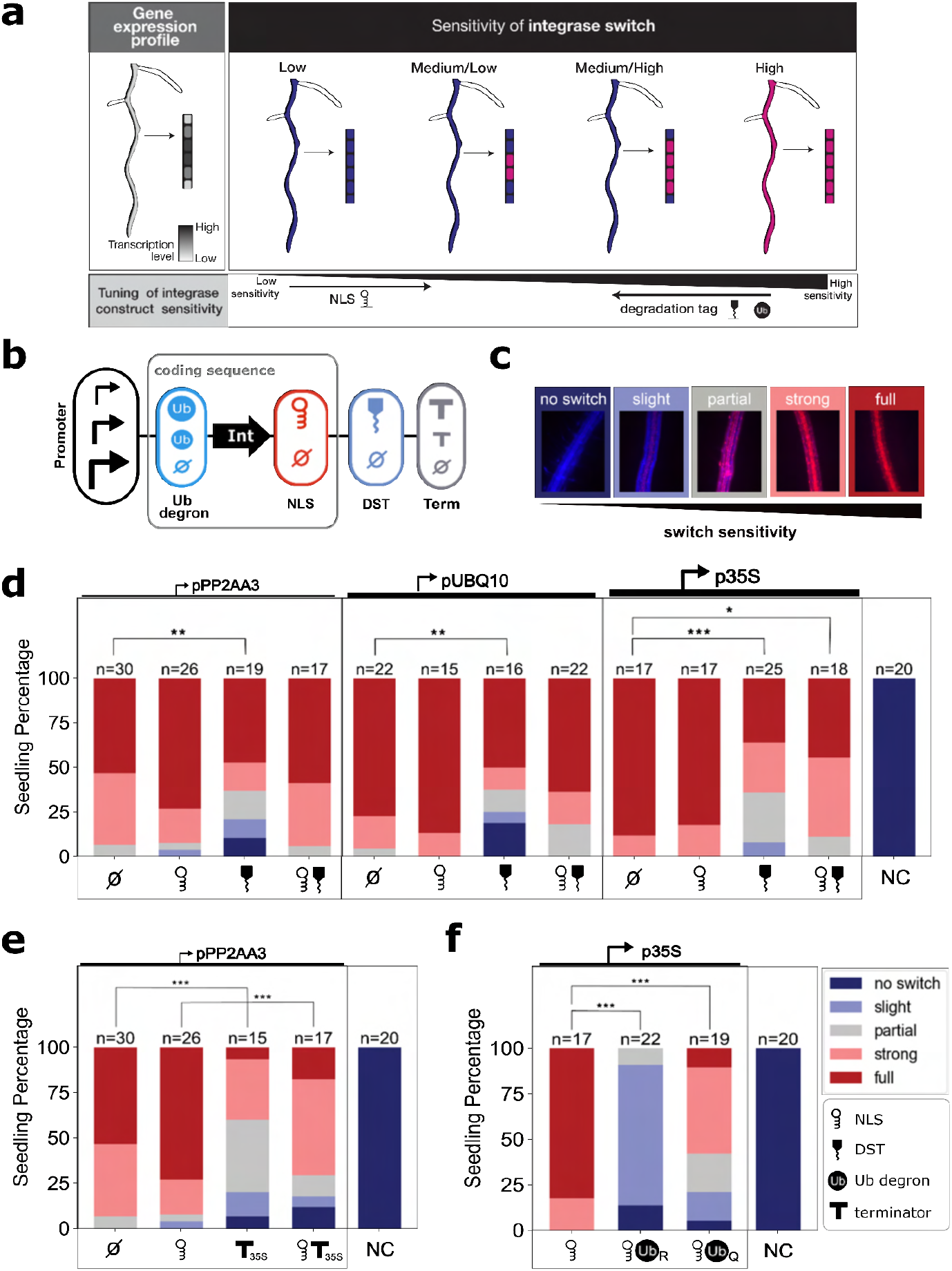
Methods for tuning integrase switch sensitivity. (**a)** For the gene expression profile of a given gene, a low sensitivity integrase switch will result in little or no switching in any cells while a high sensitivity switch will result in complete switching even in cells with relatively low expression of the gene. Different sensitivities of the integrase switch can lead to switches occurring at different levels of transcription. This sensitivity must be tuned to achieve the desired specificity for a given gene expression profile. (**b)** The integrase tuning constructs consist of a constitutive promoter controlling the integrase expression with tuning add-ons including a nuclear localization signal (NLS), RNA destabilization tag (DST), ubiquitin (Ub) degrons, and varied terminator. **(c)** The level of sensitivity is sorted into one of five categories, evaluated based on the level of mScarlet compared to mTurquoise fluorescence (Fig. S5). **(d)** Tuning results using 3 constitutive promoters (pPP2AA3, pUBQ10, p35S) with NLS, DST, and both. The negative control (NC) is the target line without any transformed integrase construct (**e, f**) Additional tuning methods include varied terminator (**e**) and Ub degrons (**f**). For each construct, seedlings were categorized into one of five classes, the percentage of seedlings in each category were plotted in a bar plot with the number of seedlings tested mentioned at the top of the bar. To evaluate statistical significance, each switching category was assigned a number from 1 through 5 (1=no switch, 5=full switch). Significance was determined using analysis of variance (ANOVA) with a post-hoc Tukey’s Honest Significance Difference test (*: p<0.05, **: p<0.01, ***: p<0.001).

### A suite of tools to optimize switch sensitivity by tuning integrase activity

The integrase switch is a binary output, while gene expression is analog and conventionally defined relative to a standard “background” or “basal” level. For example, low level gene expression is often “rounded down” to be defined as off when it falls below an arbitrary threshold, and is considered specific to a developmental event when enriched above a similarly arbitrary threshold. To be able to record events marked by different promoters, each with their own relative levels of “off” and “on”, we needed to be able to tune the sensitivity of the integrase switch (e.g., at what level of promoter expression the integrase switch is activated; Fig. 3a). While there is a rich literature of characterizing modular modifications for tuning protein activity in other systems, there are relatively few such parts available for plant synthetic biologists. We decided to characterize modifications that were predicted to work at the transcriptional, translational, and post-translational level (Fig. 3b).

We tested our tuning methods by expressing the integrase construct constitutively and observing the resulting level of switching in the roots of T1 seedlings. While in theory the constitutively expressed integrase should cause every cell to behave in the same manner, stochasticity in transcription, translation, and integrase activity results in cell-to-cell variation in the precise timing when switching occurs. This variation makes it possible to use the level of switching at a given time point as a performance metric that serves as a proxy for integrase activity. To capture the range of variation observed, each seedling was assigned to one of five classes, capturing the relative level of mTurq to mScarlet observed (Fig. 3c, Fig. S5). The classes ranged from no switching (mTurq only) to full switching (mScarlet only).

To mimic integrase expression under developmental promoters of different strengths and to capture the impact of transcriptional control modifications, we used three constitutive promoters of increasing strengths: pPP2AA3, pUBQ10, and p35S (Fig. 3d). We observed subtle differences in the switching behavior among the various promoters with p35S-driven integrase lines showing the highest percentage of seedlings in the full-switch class (Fig. 3d). As a further test of transcriptional control and in recognition of recent work documenting the striking impact of terminator sequences on gene expression^42^, we switched the UBQ1 terminator for one of our promoters, pPP2AA3, to the 35S terminator. We found that the constructs with t35S showed a decreased switch sensitivity compared to those with tUBQ1 (p<0.001) (Fig. 3e). This result further highlights the importance of promoter-terminator interactions, which could involve loop formation or the preferential localization of transcription factors to different terminator regions^43, 44^.

For post-transcriptional modifications, we studied the impacts of an SV40 T antigen-derived nuclear localization signal (NLS) (^45^) predicted to increase integrase activity^46^ and an RNA destabilization tag (DST) from SMALL AUXIN UP-REGULATED RNA (SAUR) genes^47^ predicted to decrease activity (Fig. 3b). For pPP2AA3, the addition of the NLS appeared to increase the proportion of fully switched seedlings when compared to the construct with the integrase alone, although the difference was not statistically significant (p=0.15) (Fig. 3d). For the stronger promoters pUBQ10 and p35S, the addition of the NLS did not significantly affect the switching threshold, which was likely already at a maximum level. The addition of the DST significantly decreased the switch sensitivity for all three promoters (pPP2AA3: p<0.01, pUBQ10, p35S: p<0.001) (Fig. 3d). In all cases, the addition of the DST increased the proportion of seedlings in the weaker switch categories and, in the case of pUBQ10 and p35S, reduced the proportion of fully switched seedlings.

As a final option for post-translational tuning, we tested two ubiquitin (Ub)-based protein destabilization tags (Fig. 3e). These tags work by exposing an N-terminal residue which triggers degradation of the protein by ubiquitin ligases^48^. Previously characterized in *Saccharomyces cerevisiae* (yeast), the N-end rule states that the identity of the N terminal residue determines the half-life of the protein, thus different Ub degrons confer varying levels of instability^49^. We chose a Ub-Arginine (UbR) and a Ub-Glutamine (UbQ) degron to test as in yeast they had a strong or modest impact on protein turnover, respectively^50^. While the N-degron pathway has been characterized in plants^51^, it has not been used in synthetic circuits *in planta* to tune protein levels. Consistent with the yeast results, we found that both degrons significantly increased the threshold for the integrase switch when compared to the integrase alone (p<0.001 for both comparisons), with UbR acting more strongly than UbQ.

As transient expression in *N. benthamiana* is a favorite testbed for plant synthetic biology applications, we wanted to know whether this toolbox of tuning strategies would be useful in that context as well. In addition to the pPP2AA3, p35S, and pUBQ10 promoters, we also tested the collection of tuning options with pARF19, another weak promoter known to be expressed in *N. benthamiana* leaves^52^. The NLS had a similar effect as in *Arabidopsis,* increasing the switch sensitivity for the integrase expressed under pPP2AA3 and pARF19 but not pUBQ10 or p35S (Fig. S6). Unlike in *Arabidopsis*, the DST did not have a significant effect on switching in *N. benthamiana (*Fig. S6). The 35S terminator with the NLS significantly increased switching and without the NLS tag increased switching with approaching statistical significance (p=0.17) as compared to the UBQ1 terminator in *N. benthamiana* (Fig. S6). This is in contrast to our findings in *Arabidopsis.* Additionally, the effect of the Ub degrons was quite different from what was observed in *Arabidopsis (*Fig. S6). UbR, which drastically reduced switching in *Arabidopsis,* did not significantly affect the switching in *N. benthamiana.* Even more surprisingly, UbQ which conferred a more modest, but significant reduction in switch sensitivity in *Arabidopsis*, increased switching in *N. benthamiana*. These differences have practical implications for optimizing synthetic devices, but also point to potentially fundamental differences in N-end rule dynamics and control between the two plants.

### Engineering robust integrase switches with developmental promoters

We next wanted to test the impact of the tuning modifications for developmental promoters, and focused on the impact of the NLS and DST in combination with pGATA23 and pARF19 constructs. As expected, the NLS increased the proportion of seedlings showing non-exclusive switching from less than 10% to above 86% (Fig. 4a). Conversely, the addition of the DST led to the absence of non-exclusive switching, and a slight increase in seedlings with no observed switch (from 5% no-switch in pGATA23 alone to 9% with DST; from 9% no-switch in pARF19 alone to 10% with DST) (Fig. 4a). Similarly, for pLBD16, the addition of the NLS leads to non-exclusive switching in 100% of seedlings, with 97% of the seedlings fully switched to mScarlet expression in the entire seedling (as opposed to the small number of non-LR cells showing switching in the transgenics expressing unmodified integrase from pLBD16) (Fig. 4a and Fig. S3). When pLBD16 was used to drive expression of an integrase modified with the DST, no seedlings were categorized as LR-only (Fig. 4a), but there were a higher proportion of seedlings with a switch only in the LR and in a few cells in the main root (76% with DST, 47% without; Fig. S3). This trend is consistent with the DST allowing recording of which cells express the highest level of *LBD16*.

**Figure 4:**
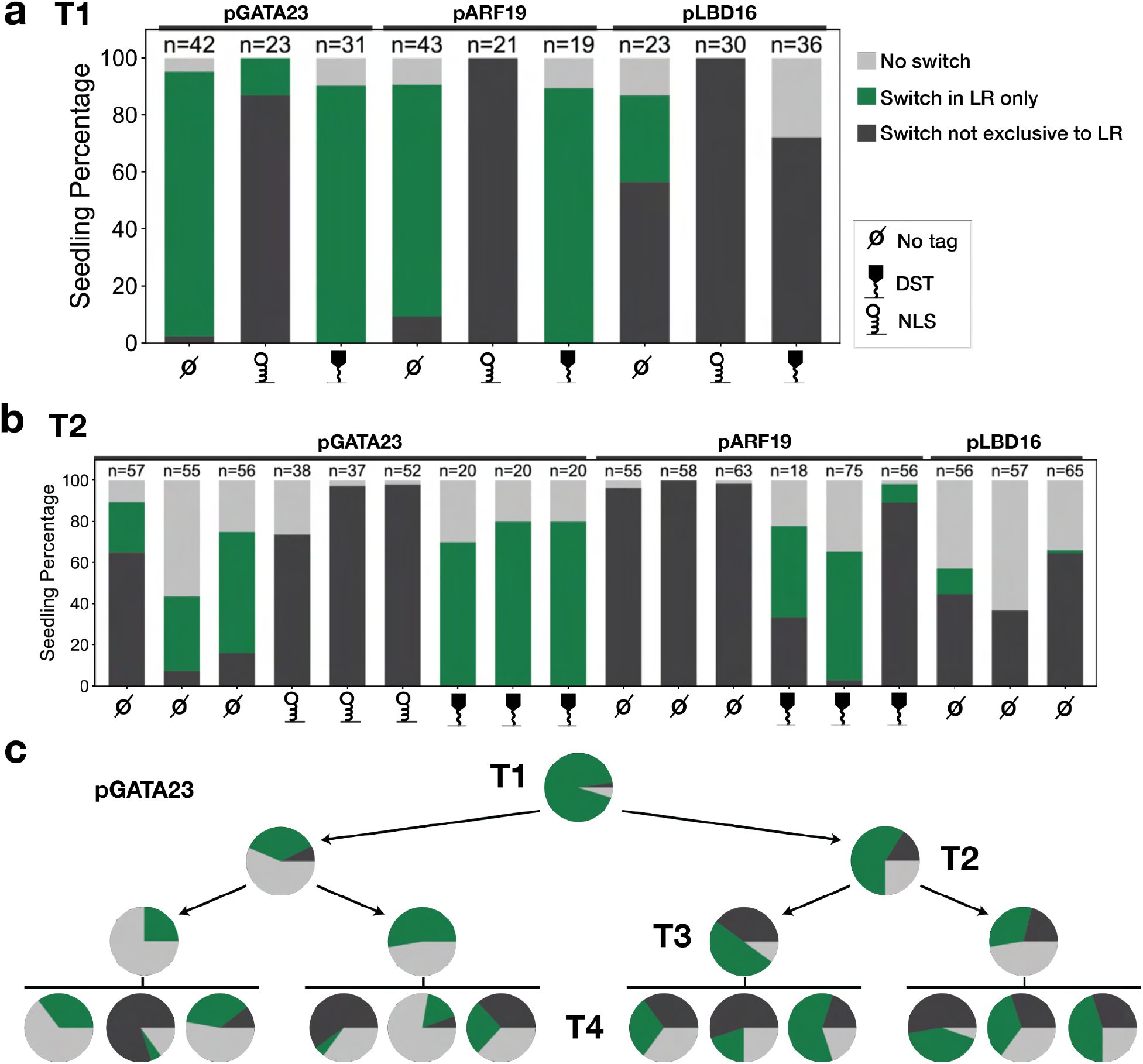
Developmental promoters with tuning tags are stable over multiple generations. **(a)** Phenotyping of T1 seedlings with constructs in PhiC31 target line, PhiC31 integrase is expressed from the indicated developmental promoters in combination with various tuning tags (legend on the right). The percentage of seedlings in each of the defined phenotypic categories is shown. (b) Phenotyping of T2 seedlings from a subset of the T1 lines represented in (a). T2 from three T1 lines per construct were characterized, all the T1 lines selected for T2 were switching only in the LR. (c) Percentage of pGATA23::PhiC31 seedlings in each phenotypic category over four generations. The pie charts for T1 and T2 are derived from the same data as displayed in (a) and (b). From each generation, three seedlings categorized as LR-only were selected for propagation.

As for the constitutive promoters, we tracked the stability of integrase switching behavior between generations. As seen previously, the ratio between switching categories changed somewhat between T1s and T2s. For pGATA23, the addition of the DST increased the stability of the phenotypic ratio of the T2 generation (Fig. 4b). We did not observe any T2 seedlings with non-exclusive switches while using the DST. The presumed increase in integrase efficiency with the NLS led to a complete absence of T2 seedlings with an LR-only switch, consistent with the T1 pattern. For pARF19, the integrase switch was no longer LR specific in T2s (Fig. 2b). The addition of the DST increased the stability of the switch in the T2 generation, leading to an LR-only switch of up to 47% of seedlings in one T1 family (Fig. 4b). We still observed a high variability between families, with some showing mostly non-exclusive switching (Fig. 4b and Fig. S7). For pLBD16, the T2s had a higher proportion of non-exclusive seedlings, although the addition of the DST narrowed the extent of mScarlet expression outside the LR (Fig. 4b and Fig. S3). We also investigated the extent to which no-switch T2s represented individuals that had lost the integrase (as T2s were not selected on antibiotic before characterization). After performing a post-characterization selection, we found that the proportion of no-switch seedlings was highly reduced (Fig. S8), meaning that we are likely underreporting the stability of the lines in T2s.

To analyze later generations, we followed a T1 carrying pGATA23::PhiC31 over four generations, propagating two LR-only seedlings at each generation. In the fourth generation, we obtained in median 32.5% (ranging from 5 to 60%) of seedlings with an LR-only phenotype (Fig. 4c). While there was clear line-to-line variability and some loss of phenotypic robustness, we could find many lines where the integrase-based recorder was still working well even in the T4 generation. The addition of the DST appeared to further stabilize the recorder function, as 65 to 100% of T3 seedlings were LR-only and there were no seedlings with non-exclusive switching phenotypes (Fig. S7c).

In addition to wanting performance stability across generations, we also wanted to make sure that an integrase-based recorder would faithfully record the spatiotemporal pattern of developmental gene expression, as there is an inherent lag between the induction of a promoter of interest and the time when the switched target reporter is detectable. To test whether this time gap was relevant to the timescales of lateral root development, we compared the expression pattern of our genes of interest using a traditional transcriptional reporter with the expression pattern of the integrase switch system driven by the same promoter. While our initial characterization shown in Fig. 2 revealed the overall pattern of integrase-based recorder activity, for these comparisons we focused our attention at the earliest stage of lateral root development. Onset of expression for transcriptional reporters and integrase constructs appeared essentially identical (Fig. 5, Dataset 4), indicating that the integrase system records the spatiotemporal pattern of gene expression with no significant delay. Beyond allowing for heritable gene expression in all daughter cells, an additional benefit of the integrase system was amplification of the developmental promoter signals. This was most obvious with the weakest promoter, pLBD16. By the same logic, the integrase system could be of great use for any application requiring normalization of output levels from multiple promoters or across multiple input signals.

**Figure 5:**
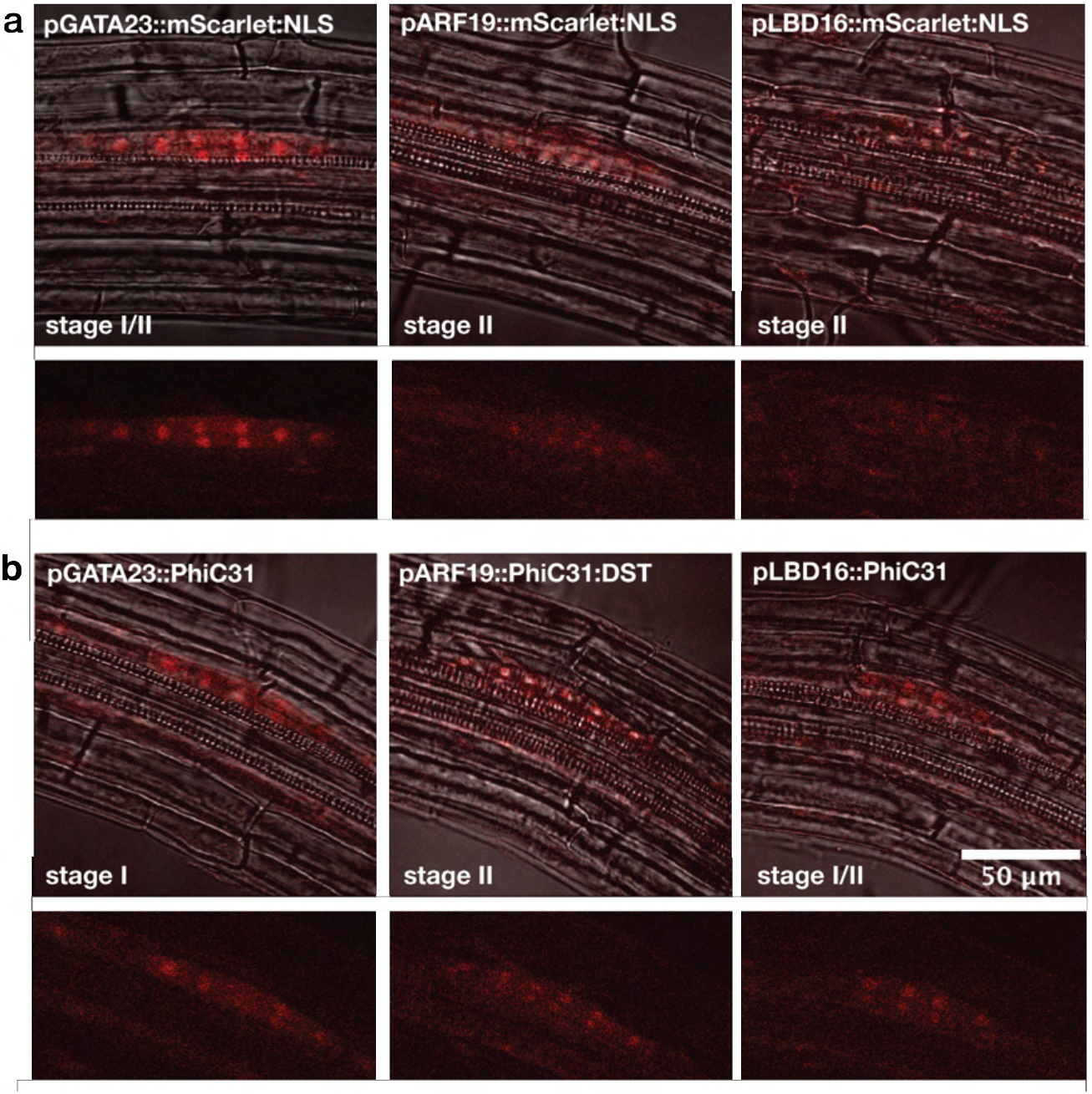
Confocal imaging of transcriptional reporters and integrase-based recorder in early-stage lateral roots. (a, b) Overlays of brightfield and red fluorescence channels from a single frame are shown above an image of the red fluorescence channel alone. For each image, the developmental stage of the lateral root primordium is indicated. (a) Transcriptional reporter lines, composed of the promoter of interest driving expression of mScarlet fused to an NLS. (b) PhiC31 integrase-based recorder lines with any modifications indicated in each panel.

### Increasing the potential applications of the integrase-based recorder

Another challenge with the integrase-based recorder is that many cellular events of potential interest may not have well-characterized promoters associated with them, or may rely on promoters that are activated in multiple cell types or conditions. For example, any promoter active in the embryo could trigger the switch of the integrase target, making any subsequent recording impossible. To overcome this limitation, we built additional tools that allow induction of the integrase at a user-determined time in development.

The first of these tools is the split intein integrase system which has been characterized *in vitro* ^53^. Inteins are sequences that trigger autocatalytic splicing, making it possible to reconstitute proteins from fragments expressed from two separate constructs^54^, a technique that has been used previously in plants^55^. In the split intein integrase system we applied here, the PhiC31 integrase is split into two extein domains: the N-terminal sequence fused to the intein N-term: Npu DnaEN and the PhiC31 C-terminal sequence fused to the intein C-term: Ssp DnaEC (Fig. 6a). Expression of the two components in a single cell triggers post-transcriptional trans-splicing, generating a fully functional PhiC31 integrase. We tested the split-intein integrase system in *Arabidopsis* with the PhiC31 integrase using strong constitutive promoters for the expression of the two components: pUBQ10 for the N-term, and p35S for the C-term, and found that it worked well (Fig. 6b). We compared the level of integrase switch from this construct to the full integrase under the control of each of the constituent promoters, and found that the split-intein system led to a decrease in switch efficiency. While 90% of split-intein T1 seedlings showed some level of switching, no full switch was observed. In contrast, when the full integrase was expressed under the control of either pUBQ10 or p35S, 75% or more of the seedlings were fully switched. This is consistent with reports that the trans-splicing approach delays the integrase switch in *E. coli*^53^.

**Figure 6:**
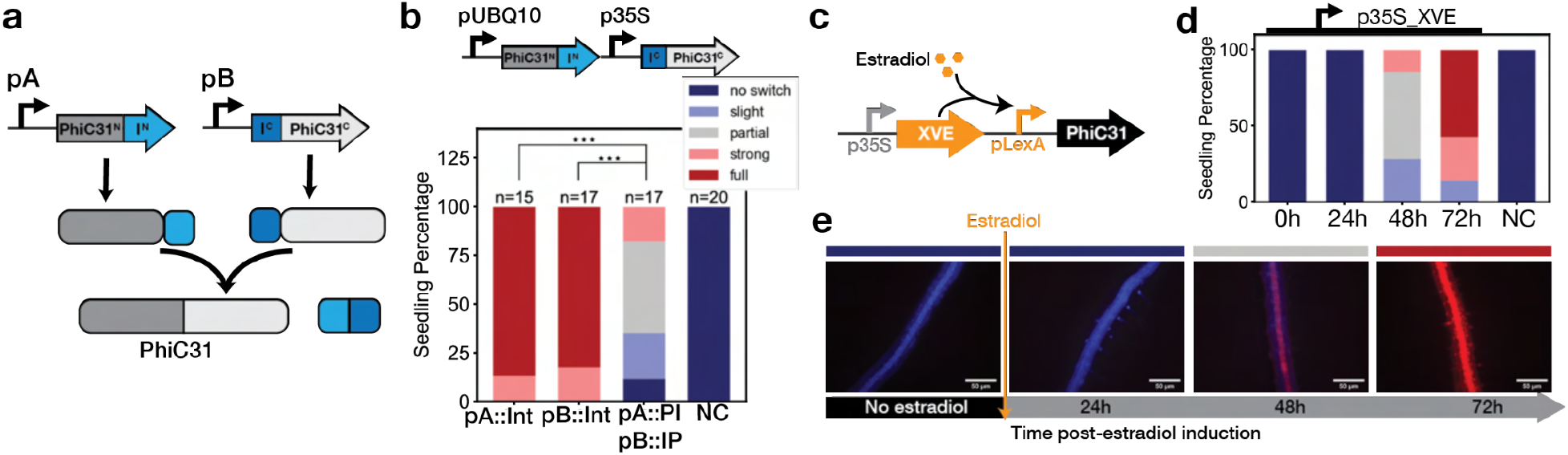
Split-intein and inducible promoters as additional tools to tune and induce the integrase switch. (a) Schematic of the split-intein integrase system. The system is composed of two constructs: (i) promoter A driving the N-terminal half of the integrase (PhiC31^N^, dark gray) fused to the N-terminal portion of the intein protein (I^N^: Npu DnaE^N^, light blue) and (ii) promoter B driving the C-terminal portion of the intein (I^C^: Ssp DnaE^C^, dark blue) fused to C-terminal half of the integrase (PhiC31^C^, light gray). When the two constructs are expressed, the inteins autocatalyze trans-splicing, covalently joining the two parts of PhiC31. (b) The split-intein system reduces the efficiency of the integrase switch. Following the nomenclature of (a), pUBQ10=promoter A and p35=promoter B. The integrase switch efficiency of the split-intein system is compared with the full integrase expressed with promoter A alone (pUBQ10:: PhiC31) and with promoter B alone (p35S::PhiC31). NC corresponds to the target line without integrase. T1 seedling phenotypes are determined with fluorescent microscopy images and categorized from no switch to full switch. The data were tested for significance using an ANOVA and post-hoc Tukey’s HSD test (*: p<0.05, **: p<0.01, ***: p<0.001). (c) The estradiol inducible integrase construct is composed of p35S::XVE (transcriptional activator composed of a DNA-binding domain of LexA, the transcription activation domain of VP16, and the regulatory region of the human estrogen receptor;^57^) and pLexA-minimal 35S driving expression of PhiC31. (d) Characterization of the estradiol inducible integrase construct shows induction as early as 48 h after treatment. T1 Seedlings with the estradiol integrase construct in the PhiC31 target line were characterized just before estradiol treatment and every 24 hours following. The bar graph represents the percentage of seedlings with a given level of switching (classes are color coded as in (b)), n=14 seedlings. (e) Representative images of a seedling at the specified time point relative to estradiol induction.

The split-intein integrase system could be used to induce the integrase recording system at a specific stage of seedling development, thereby avoiding recording at earlier stages. This would be done by placing one component under the control of a developmental promoter and the other under the control of an inducible promoter. We could then activate the integrase system through the inducible promoter at the beginning of an experiment to record the expression of genes only after a specific time point, reducing issues with genes expressed in embryonic tissues. As a proof-of-principle for this design, we used the heterologous estrogen inducible system^56, 57^ to drive integrase expression. Before estradiol induction, we did not observe any switch, confirming that the estradiol system had an undetectable level of background activity (Fig. 6c). After induction, we analyzed seedlings every 24 hours and observed the earliest signs of switching at 48h with more than 50% of seedlings fully switched by 72h. This timing fits well with reports that estradiol induction of a reporter peaks at 24h^57^, and would suggest that it takes approximately 24 hours after promoter activation for the integrase to become active, mediate the switch and then allow expression and maturation of mScarlet. We also characterized our inducible integrase construct in T2s and confirmed that the baseline of expression without inducer is low enough to prevent any integrase switching in subsequent generations (Fig. S9).

## Discussion

Integrase-based recorders of gene expression have a number of advantages over current methods of tracking transcription in individual cells. Among the most prominent of these is that early events can be read-out much later in development, and, in the designs presented here, there is no need to disrupt the spatial relationship between cells. We have added a suite of characterized parts to help synthetic biologists build devices in *Arabidopsis*. In addition to the PhiC31 and Bxb1 integrases and cognate targets, we built and tested tuning modifiers like RNA and protein destabilization tags and a split-intein control module. This entire suite of standardized tools can be directly implemented in any system where fine control of protein levels is needed to optimize performance. We also provided proof-of-principle that integrase-based recorders can be used to capture the history of gene expression at specific times and spaces during plant development. Importantly, we also found that switches functioned robustly over multiple generations. Additionally, we observed differences in the effect of tuning parts between *Arabidopsis* stable lines and *N. benthamiana* transient assays, adding another note of caution in developing synthetic devices for use in multiple plants.

The integrase-based recording system characterized here can be readily adapted to the tracing activity of other promoters, including those expressed in other tissues and developmental processes. Because the integrase acts as a signal amplifier, the integrase system could be of interest in following expression of any genes that are difficult or impossible to observe with traditional reporters. Additionally, the integrase system could be used to record the expression of genes in situations where live imaging is not available. For example, while many labs have at least some access to fluorescent microscopy, most do not have access to sophisticated live-imaging set-ups. There are also conditions, such as roots growing in natural soil conditions, where it would be highly advantageous to read-out early expression events much later in development. Moreover, in situations where imaging is not compatible with other protocols (e.g., some fixation techniques), it is also possible to detect the state of the integrase targets used here by sequencing.

Additional synthetic devices should now be accessible working from the toolbox described here. For example, one challenge in producing a developmental recorder is that many promoters of interest are expressed at multiple points in development. One solution would be to combine our inducible integrase and split-intein integrase system, where one part of the integrase is under the control of the externally inducible promoter and the other is expressed from the developmental promoter of interest. Another use of the split-intein integrase would be to use it as an AND gate by placing the two split-intein components under the control of two promoters from genes of interest. This will allow the recording of when and in which cells two different genes are simultaneously expressed. By using both PhiC31 and Bxb1 integrases, a history-dependent tracker could be constructed with the capacity to record on a single cell level if, and in what order, two genes are expressed. A similar design has been shown previously to work in bacteria^19^.

In addition to contributing to our understanding of existing organisms, integrase-based devices can also enable engineering of novel forms or functions by driving expression of genes other than reporters. For example, integrase switches could be used to induce the expression of a toxic gene under certain conditions. Cre recombinase, a tyrosine integrase, has already been used to generate homozygous fertilization-defective mutants in plants^58^, and to activate a large-tumor antigen in mice^59, 60^. A particularly exciting application to imagine is to replace reporter genes in integrase targets with transcription factors able to initiate entire response cascades. Root development could be re-coded by implementing history-dependent synthetic signaling circuits that used integrases to activate developmental regulators, potentially helping plants survive drought or flooding.

## Supporting information

Suppdata2

Suppdata1

Suppdata5

## Acknowledgement

We thank Wesley George, Eric Yang, Dr. Alexander Leydon, as well as other members of the Nemhauser, Imaizumi, and Steinbrenner groups, for feedback and discussions. We thank Eric Yang for developing and gifting the pPP2AA3 promoter; Dr. Jennifer Brophy for sharing the RFP injection efficiency control; and members of the Orzáez lab for useful discussions, as well as the *N. benthamiana* PhiC31 target line and the YFP to Luc PhiC31 target plasmid.

## Online methods

### Construction of plasmids

Our cloning strategy was based on Golden Gate assembly using appropriate spacers (Fig S10) ^61^ and BsaI-HFv2 (NEB) as the restriction enzyme. Candidate promoter sequences (ARF7: AT5G20730, ARF19: AT1G19220, LBD16: AT2G42430, GATA23: AT5G26930) were amplified from Col-0 genomic DNA to add specific Golden Gate spacers. After gel purification, each level0 promoter sequence was cloned using a Zero Blunt PCR Cloning Kit (ThermoFisher Scientific). The PhiC31 integrase sequence was a gift from the Orzeaz lab. Bxb1 and Tp901 sequences were a gift from the Bonnet lab. Integrases were amplified using primers with golden gate compatible spacers to generate level0 integrase parts (primer list available in Dataset 1). Constitutive plant promoters and terminators were purchased from Addgene as part of the MoClo Toolbox for Plants ^61^. Some level0 parts were ordered from Twist Bioscience: a mutated version of the pPP2AA3 promoter without BsaI sites, the DST, the Ub-tags, the mTurq-tUBQ10 level0 construct for target construction. The mScarlet-tRBCs level0 construct was amplified from a transcriptional reporter from ^10^. Other level0 fragments were ordered from IDT as Gblocks: the codon optimized Tp901 integrase sequence, the two split-intein PhC31 constructs, and the integrase target sequences without promoters. For the integrase target level0 sequences, the pUBQ10 promoter was added by Golden Gates using BbsI sites.

Construction of constitutive and lateral root specific level 1 integrase constructs was performed via Golden Gate reaction in the modified pGreenII-Hygr vector containing compatible Golden Gate sites ^62^. Construction of integrase targets was performed with the same methods in a modified pGreenII-Kan vector.

Construction of level 2 integrase constructs, such as the split-intein system construct, was performed by amplifying completed level 1 integrase constructs using primers with golden gate compatible spacers, then performing Golden Gate reactions in the modified pGreenII-Hygr vector containing compatible Golden Gate sites.

Construction of promoter reporters was performed by assembling through Golden gate reaction the mScarlet with NLS, tRBCs terminator, and promoter in the modified pGreenII-Hygr vector as in ^10^.

Details on constructs and primers can be found in Dataset 1. The sequences of all level1 constructs can be found in Dataset 5.

Enzymes for Golden Gate assembly were purchased from New England Biolabs (NEB, Ipswich, MA, USA). PCR was performed using 2X Q5 PCR master mix (NEB) and GoTaq master mix for colony PCR (Promega, Madison, WI, USA). Primers were purchased from IDT (Louvain, Belgium), and DNA fragments from Twist Bioscience or IDT. Plasmid extraction and DNA purification were performed using Monarch kits (NEB). Sequences were verified with Sanger sequencing by Azenta Life Sciences (Seattle, USA). Chemically-competent cultures of the *E. coli* strain DH5alphaZ1 (laciq, PN25-tetR, SpR, deoR, supE44, Delta(lacZYA-argFV169), Phi80 lacZDeltaM15, hsdR17(rK−, mK+), recA1, endA1, gyrA96, thi-1, relA1) were transformed with plasmid constructs containing kanamycin resistance. Transformed *E. coli* was grown in LB media (LB broth, Miller) with kanamycin (Millipore Sigma, 50 ug/mL).

### Plant growth conditions

*Arabidopsis* seedlings were sown in 0.5 X Linsmaier and Skoog nutrient medium (LS) (Caisson Laboratories) and 0.8% w/v agar, stratified at 4°C for 2 days and grown in constant light at 22°C. Phyto agar (PlantMedia/bioWORLD) was used when imaging seedlings and Bacto agar (ThermoFisher) otherwise.

### Construction and selection of transgenic *Arabidopsis* lines

*Agrobacterium tumefaciens* strain GV3101 was transformed by electroporation, and subsequently grown in LB media with rifampin (Millipore Sigma, 50 ug/mL), gentamicin (Millipore Sigma, 50 ug/mL), any antibiotics carried on the specific plasmid(s), most often kanamycin (Millipore Sigma, 50 ug/mL).

The floral dip method ^63^ was used to generate integrase target lines in Col-0, and then used to introduce each integrase construct into these established target lines.

For T1 selection: 120 mg of T1 seeds (∼2000 seeds) were sterilized using 70% ethanol and 0.05% Triton-X-100 and then washed using 95% ethanol. Seeds were resuspended in 0.1% agarose and spread onto 0.5X LS Bacto selection plates, using 25 ug/mL of kanamycin for target lines and 25 ug/mL kanamycin and 25 ug/mL hygromycin for lines with both the integrase and the target. The plates were stratified at 4°C for 48 hrs then light pulsed for 6 hrs and covered for 48 hrs ^64^. They were then grown for 4-5 days. To select transformants, tall seedlings with long roots and a vibrant green color were picked from the selection plate with sterilized tweezers and transferred to a new 0.5X LS Phyto agar plate for characterization.

### Characterization of integrase switch in *Arabidopsis* transgenic lines

T1 seedlings for each line were grown 4-5 days after transformant selection. Each selected seedling was imaged at 10X magnification using an epifluorescence microscope (Leica Biosystems, model: DMI 3000) using the RFP (exposure 500 ms, gain 1.6) and CFP (exposure 300 ms, gain 1.6) channels. Selected T1 seedlings were then transferred to soil, and at maturation T2 seeds were selected. For later generations, seedlings were sterilized similarly as for T1s, stratified, plated on an LS agar plate, grown for 4-5 days, and characterized using the epifluorescence microscope as for T1.

For the target lines, the seedlings with the highest level of mTurq expression were selected and transferred to soil to generate T2 seeds. The brightest among these lines was maintained as the target line for each integrase, and used for all later transformations of integrase constructs. For the constitutive integrase constructs in a target line, around 20-30 T1 seedlings were analyzed per construct. Each seedling was categorized into one of five classes as seen in Fig. S5 based on the level of switching. Representative images for each category were taken using the RFP and CFP channels and merged for final images. For each construct, the percentage of seedlings in each category were plotted in a bar plot with the number of seedlings tested mentioned at the top of the bar. To evaluate statistical significance, each switching category was assigned a number from 1 through 5 (1=no switch, 5=full switch). Significance was determined using analysis of variance (ANOVA) with a post-hoc Tukey’s Honest Significance Difference test.

For the YFP to Luciferase PhiC31 target, the target line and the target with pPP2AA3-PhiC31 construct were characterized. T2 seedlings from target lines with and without integrase were grown on LS plates, 7 days old seedlings were imaged with Azure c600 Gel imaging system for YFP fluorescence. Then, 100μM of Luciferin were sprayed on seedling, after one hour kept in the dark, seedlings were imaged using NightOwl LB 983 *in vivo* imager with an exposure time of 10min.

For the developmental promoter integrase constructs in PhiC31 target line, at least 20 T1 seedlings were analyzed per construct. Each seedling was categorized into one of 3 classes based on specificity of switching (LR-only, non-exclusive to LR, no-switch). Representative images for each construct were taken using the RFP and CFP channels and merged for final images. A selected number of T1 seedlings with LR-only switch were transplanted to soil to characterize the T2 generation. For each T1 line, 20 T2 seedlings were characterized in an identical way than for T1s, and similarly for T3 and T4 generations. For each construct, the percentage of seedlings in each of the three categories were plotted in a bar plot with the number of seedlings tested mentioned at the top of the bar.

Python data analysis script which includes statistical tests and plotting functions was run in version 3.9.1 and with the following package dependencies: pandas, scipy.stats, matplotlib.pyplot, matplotlib.colors, scikit_posthocs, and numpy. All images taken during seedling characterization were opened and processed using the imageJ program (version 1.53c). Each .tif image file contained the images of a seedling’s RFP and CFP channels. .tif files were processed through an imageJ macro to adjust the color lookup table, brightness, and contrast of each channel (RFP: Red, Min: 200, Max: 3000) (CFP: Blue, Min: 200, Max: 4000). After adjustment, the macro overlaid the two channels to create a composite image, rotated the image, added a scale bar, and flattened the image to produce our final processed images. Python and ImageJ script are available on Github (https://github.com/sguiz/Integrase_plant_paper) and in supplementary material and methods. Raw data is available in Dataset 2, additional microscope images are available in Dataset 3.

### Testing the hygromycin resistance of seedlings post characterization

To select T2 hygromycin resistant seedlings after characterization without selection, the roots of 7 days old seedlings were removed with a razor blade, and seedlings were then transferred onto 0.5X LS BactoAgar plates containing hygromycin. Seedlings were screened for root regrowth after seven days. In our extensive testing of control plants, not all hygromycin resistant seedlings are able to regrow roots after this stressful intervention, but all seedlings that grow roots are truly resistant.

### Characterization of the tuning constructs in *Nicotiana benthamiana*

Integrase target integrated *Nicotiana benthamiana* seeds were acquired from the Orzaez lab ^24^. This line has a stably integrated integrase target which switches from LUC firefly luciferase to YFP upon integrase expression. The plants were grown 25 days before injection.

*Agrobacterium*-mediated transient transformation of *N. benthamiana* was performed as per ^65^ using the *A. tumefaciens* strains GV3101. For each injection in addition to the *A. tumefaciens* with the integrase constructs, we injected an RFP injection efficiency control consisting of constitutively expressed mCherry (donated by Jennifer Brophy) and a construct containing a P19 gene silencing suppressor protein for enhanced transient transformation ^66^. Each *A. tumefaciens* strain was grown overnight in LB at 30°C, pelleted and incubated in MMA media (10 mM MgCl2, 10 mM MES pH 5.6, 100 µM acetosyringone) for 3 hours at room temperature with rotation. Strain density was normalized to an OD600 of 1.5 for each strain in the final mixture of strains before injection. For each integrase construct, the integrase strain, the RFP control, and P19 were injected together; we also injected as control the RFP control and P19 together, as well as the negative control P19 alone. Each *A. tumefaciens* solution was injected into 3-4 different leaves from separate plants. Four days later, hole punches were taken from each injected leaf at 3 locations, and the punches were placed in a 96 well plate. Plate reader measurements of YFP (excitation wavelength: 506 nm, emission wavelength: 541) and mCherry (excitation wavelength: 584 nm, emission wavelength: 610) fluorescence were taken using a Spark® Multimode Microplate Reader by Tecan. Twelve measurements were taken at different locations within the punch. Three tobacco injection replicates per construct were performed and, in each replicate, three leaves were injected. For each punch, the median of the ratio of YFP over RFP fluorescence was calculated and plotted. The box corresponds to the quartile and the median between the different punches for one construct. The data were tested for significance using an ANOVA and post-hoc Tukey’s HSD test. The tobacco injection data was plotted and statistically analyzed using an Python data analysis script. Python script is available on Github (https://github.com/sguiz/Integrase_plant_paper) and in supplementary material and methods.

### Confocal imaging of reporter and integrase lines

*Arabidopsis* transgenic reporter lines for *LBD16, ARF19*, and *GATA23* with mScarlet nuclear localized were generated as previously described. After characterization of T1 seedlings, seedlings expressing mScarlet were fixed in 4% formaldehyde using vacuum infiltration followed by ClearSee solution ^67^. Fixed and cleared seedlings were mounted on microscope slides using 50% glycerol and Parafilm edges to prevent the coverslips from pressing on the root.

For the integrase lines, for each promoter *LBD16, ARF19*, or *GATA23*, one construct showing a reliable LR-only integrase switch was selected. For each construct, two T1 lines representative of other characterized T1 lines were selected to perform the root bend essay. For each line, 20 T2 seeds of the corresponding T1 line were placed on plates following a specific pattern to avoid seedling collision after the rotation of the plate. The seeds were stratified for 120h, grown vertically for 96h at 22°C, rotated 90° while keeping the plate vertical, and grown for an additional 20h. Seedlings were fixed and mounted as previously mentioned. Imaging of the seedlings were performed using Nikon A1R HD25 laser scanning confocal microscope with 561 laser and 578-623 detector for RFP imaging. For the integrase lines, seedlings were imaged at the bend region, while for the reporter lines, seedlings were scanned to find early developed lateral roots. Imaging was processed using FIJI. For each imaging, a Z-stack was recorded. First, a maximum average of the Z-stack in the RFP channel was generated. Additionally, we selected one Z-location focusing on the LR nucleus and generated both an image of the RFP channel and the RFP and brightfield merged. The main figure uses the merged RFP/brightfield images.

### Estradiol induction time course

For estradiol induction in T1s, antibiotic selection was performed as described above. Four days after transplanting resistant seedlings onto 0.5X LS Phyto plates, the seedlings are imaged as previously described in RFP and CFP channels. Then the seedlings were transferred onto new 0.5X LS Phyto plates with 10 uM β-estradiol. Each seedling was imaged 24, 48, and 72 hours after transplanting onto estradiol and categorized into the appropriate switching category for each timepoint. Data was processed as specified above.

For estradiol induction in T2s, seeds were plated onto 0.5X LS Phyto plates, stratified for 48 hours and left to grow for 6 days. Then they were transplanted onto 10 uM estradiol 0.5X LS Phyto plates and imaged and categorized as described above.

Raw data is available in Dataset 2.

## Supplementary figures

**Fig S1:**
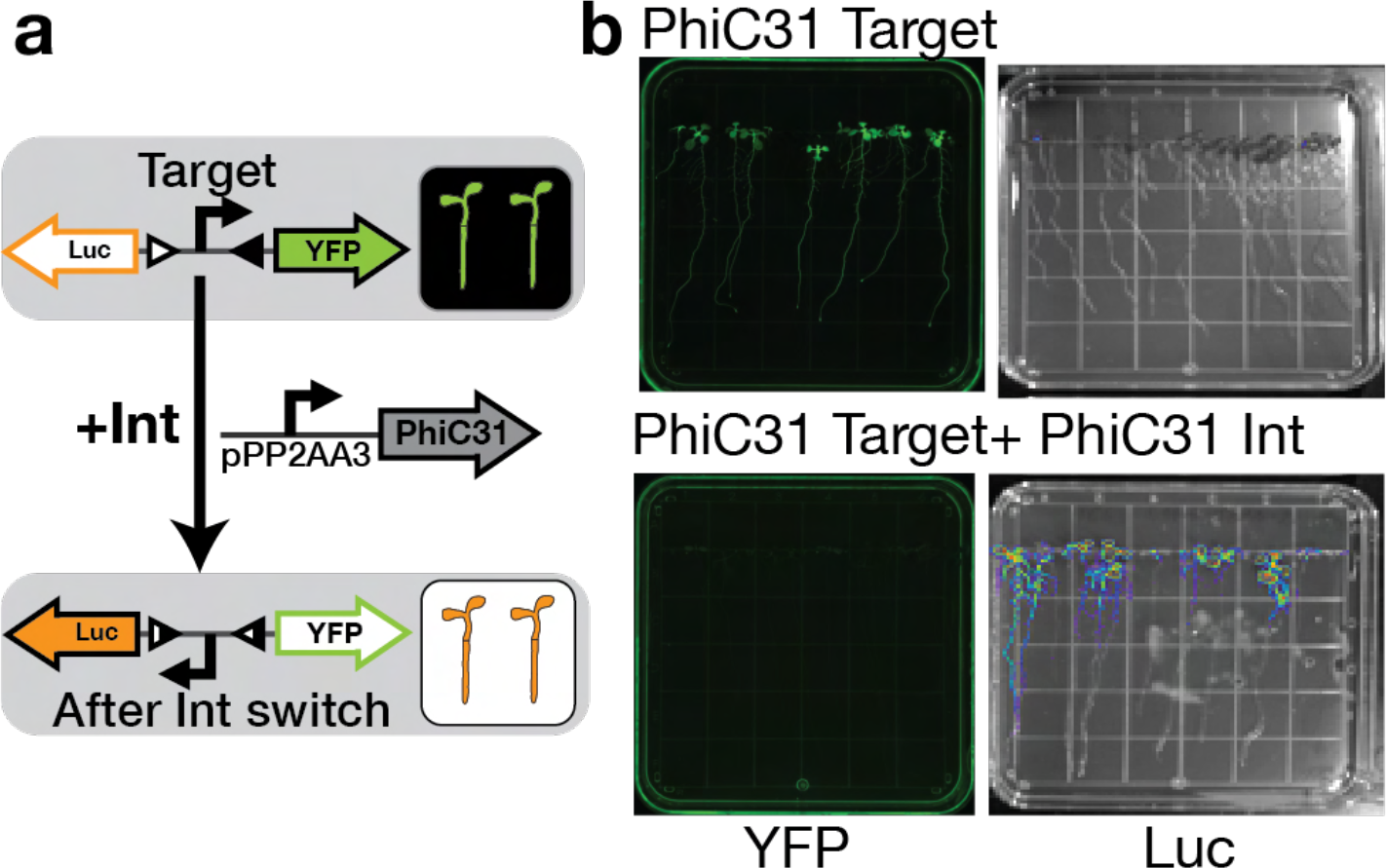
Integrase target for macroscopic analysis. (a) The PhiC31 target (from <u>(Bernabé-Orts et al.</u> <u>2020)</u>) switches from YFP to Luciferase expression, allowing to image the full seedling under gel imager for YFP or night owl after luciferin treatment. We used pPP2AA3 to drive PhiC31 integrase expression constitutively. (b) Images of T2 seedling under an Azure c600 Gel imaging system (for YFP fluorescence) (left) and NightOWL LB 983 *in vivo* imaging system (for Luciferase) after luciferin treatment (right), on the top are seedlings with target and no integrase at the bottom seedlings with target and integrase.

**Fig S2:**
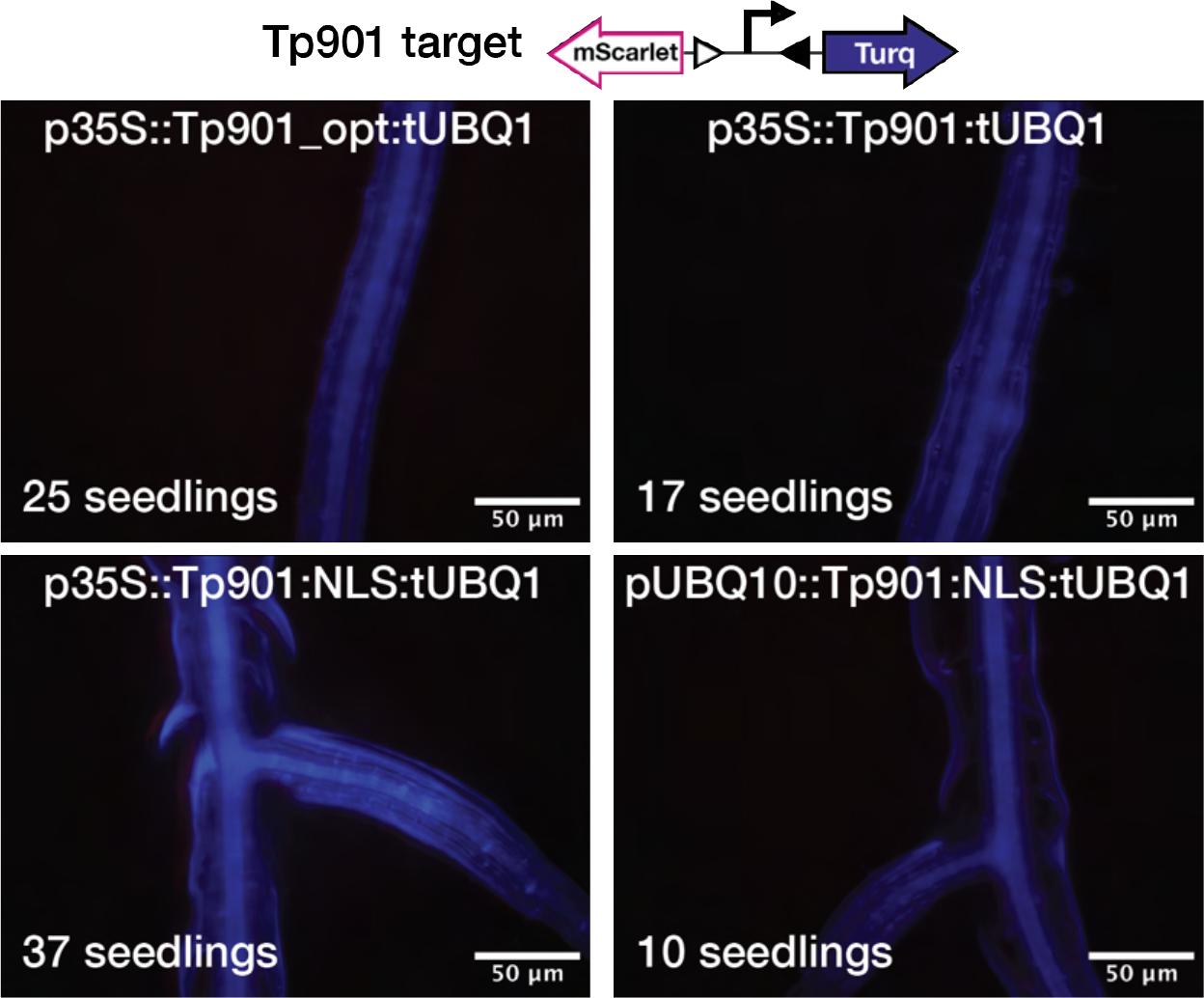
Tp901 integrase does not work well in *Arabidopsis*. Tp901 target line was transformed with Tp901 integrase constructs. If the integrase was active the target should swtch from mTurq to mScarlet expression. No mScarlet was ever detected. T1 expression patterns were analyzed by microscopy. Each image corresponds to a representative image for each integrase construct, the integrase construct is mentioned on the top and the number of seedling characterization on the bottom left. The microscopy image is an overlay of the blue and red fluorescence channel.

**Fig S3:**
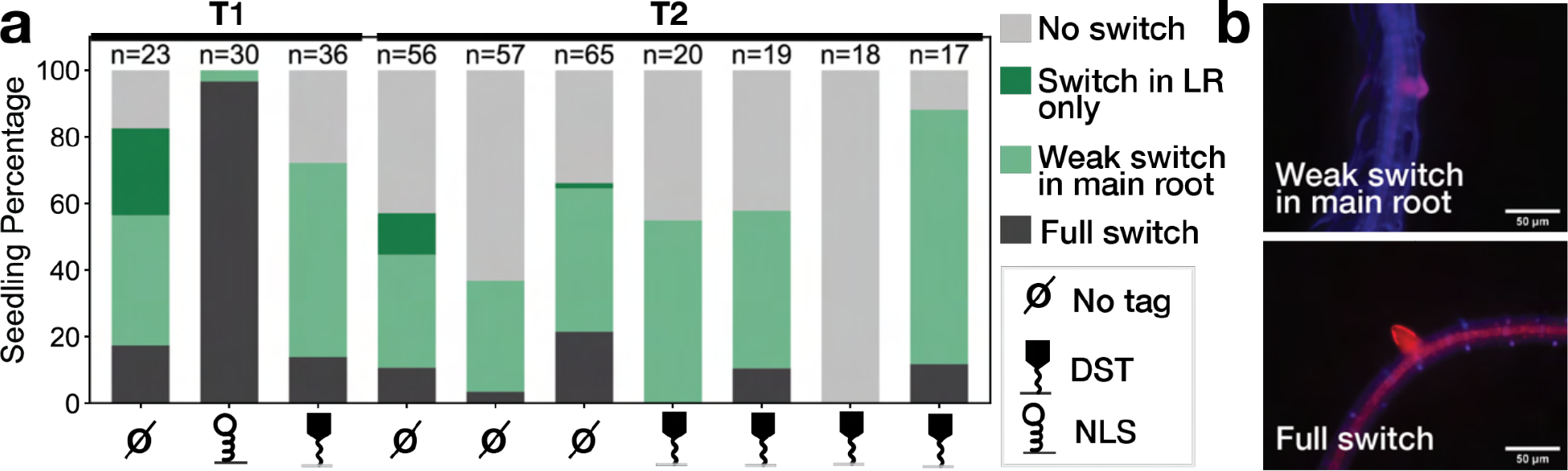
Characterization of pLBD16 integrase switch constructs. (a) Phenotyping of T1 seedlings with constructs in PhiC31 target line, constructs are PhiC31 with pLBD16, and various tuning tags (label at the bottom of the graph): no tag, DST, or NLS (legend on the right). The graph corresponds to the percentage of seedlings in each of the defined phenotype categories, such as no switch corresponding to no mScarlet expression in the root, switch in LR only: mScarlet expression only in the lateral root, weak switch in the main root: mScarlet expression in few cells in the main root (corresponding to the image in b), full switch: mScarlet expression everywhere in the root (corresponding to the image in b). The number of seedlings characterized for each construct is mentioned at the top of the bar in the graph. (b) Representative image of the seedling in the weak switch in main root and full switch phenotypic categories. Microscopy images are an overlay of the blue and red channels.

**Fig S4:**
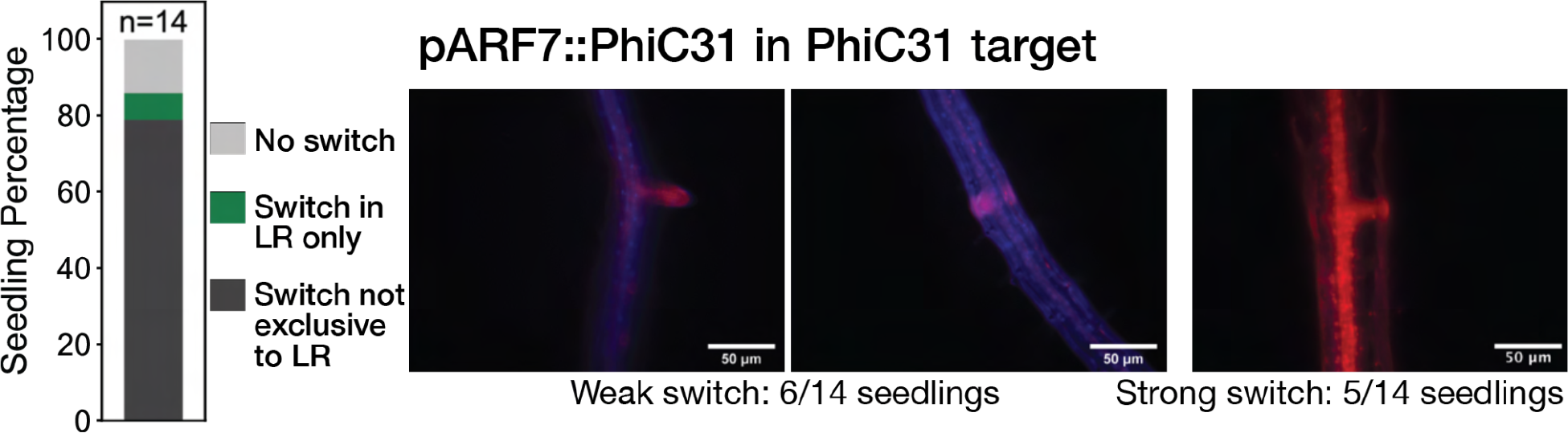
Characterization of pARF7::PhiC31 switch. 14 T1 seedlings of pARF7::PhiC31 in PhiC31 target line were characterized. On the left, the bar graph corresponds to the percentage of seedlings in each of the phenotypic categories, no switch in light gray, switch only in LR in green, and switch not exclusive to LR in dark gray. On the right are representative images of seedlings with switch not exclusive to LR, either a weak switch (6/14 seedlings) or a strong switch everywhere (5/14 seedlings).

**Fig S5:**
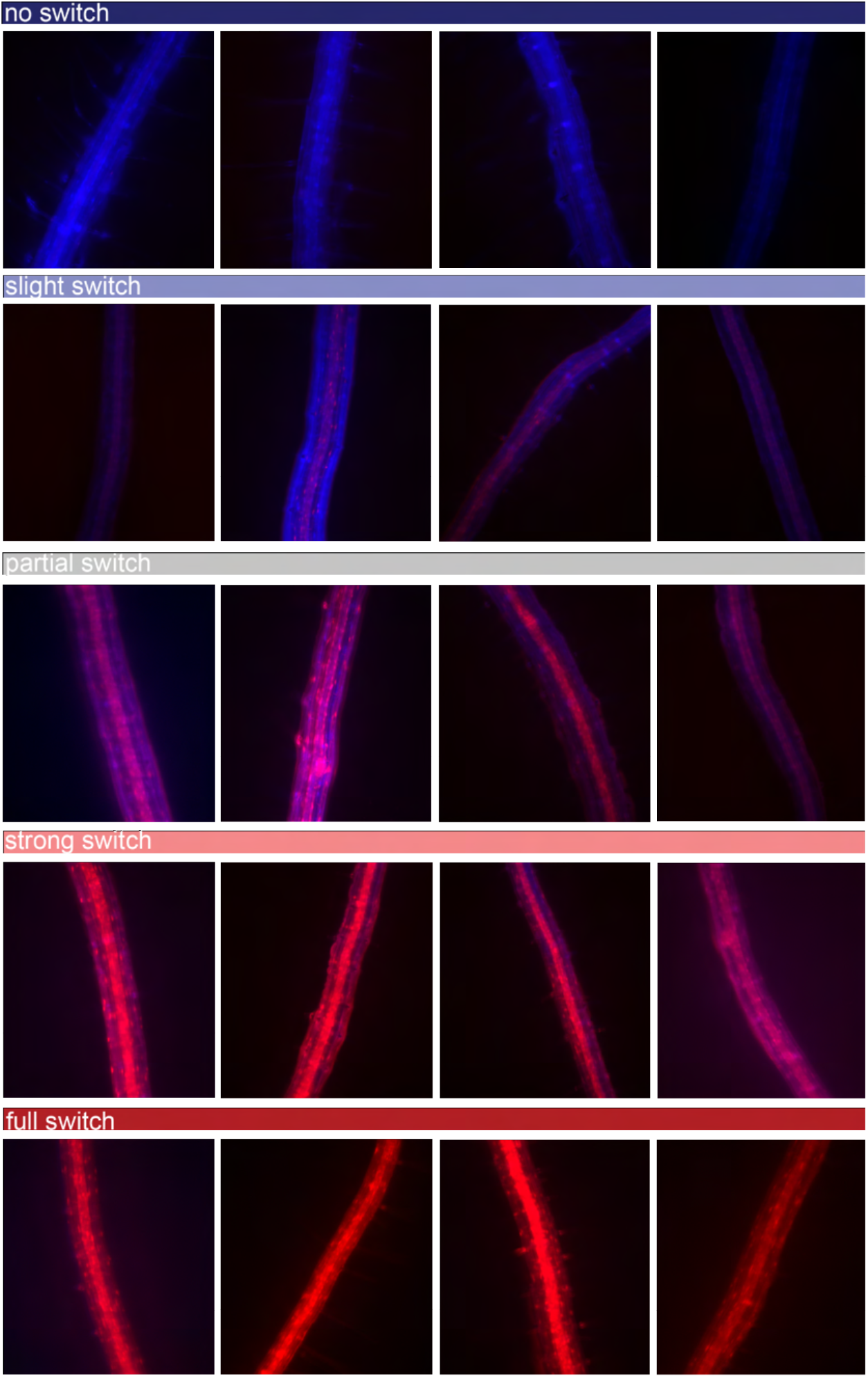
Example images of switching categories for tuning data (Figure 3). For evaluating the tuning results we sorted each seedling into one of five categories (no switch, slight switch, partial switch, strong switch, full switch). Within each category variation is present.

**Fig S6:**
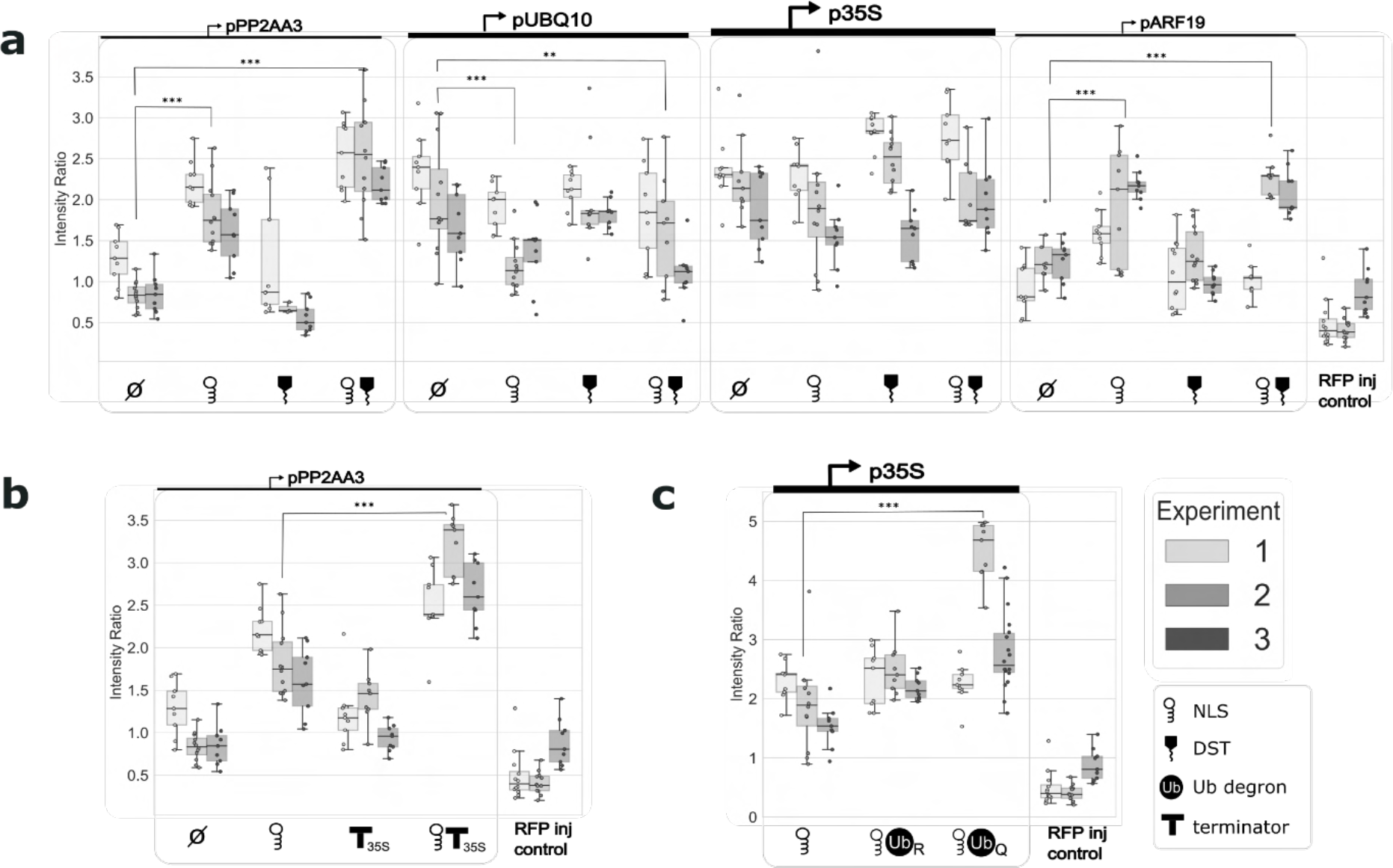
The effect of tuning modifications differ between *N. benthamiana* and *Arabidopsis*: (**a-c**) Tuning was tested in *N. benthamiana* through tobacco injection. The integrase target switches from a Luc reporter to YFP and an RFP injection control was co-injected with each construct. The metric for level of switching is the ratio of YFP to RFP. Each tuning construct was injected into 3 leaves per experiment. Three punches were taken from each leaf and the resultant fluorescence was measured with a plate reader. Each point on the boxplot represents one leaf punch. Each box represents one of three replicate experiments performed for each construct. Tuning parts tested were (**a**) NLS and DST, (**b**) Ub degron, and (**c**) varied terminator. The data were tested for significance using an ANOVA and post-hoc Tukey’s HSD test (*: p<0.05, **: p<0.01, ***: p<0.001).

**Fig S7:**
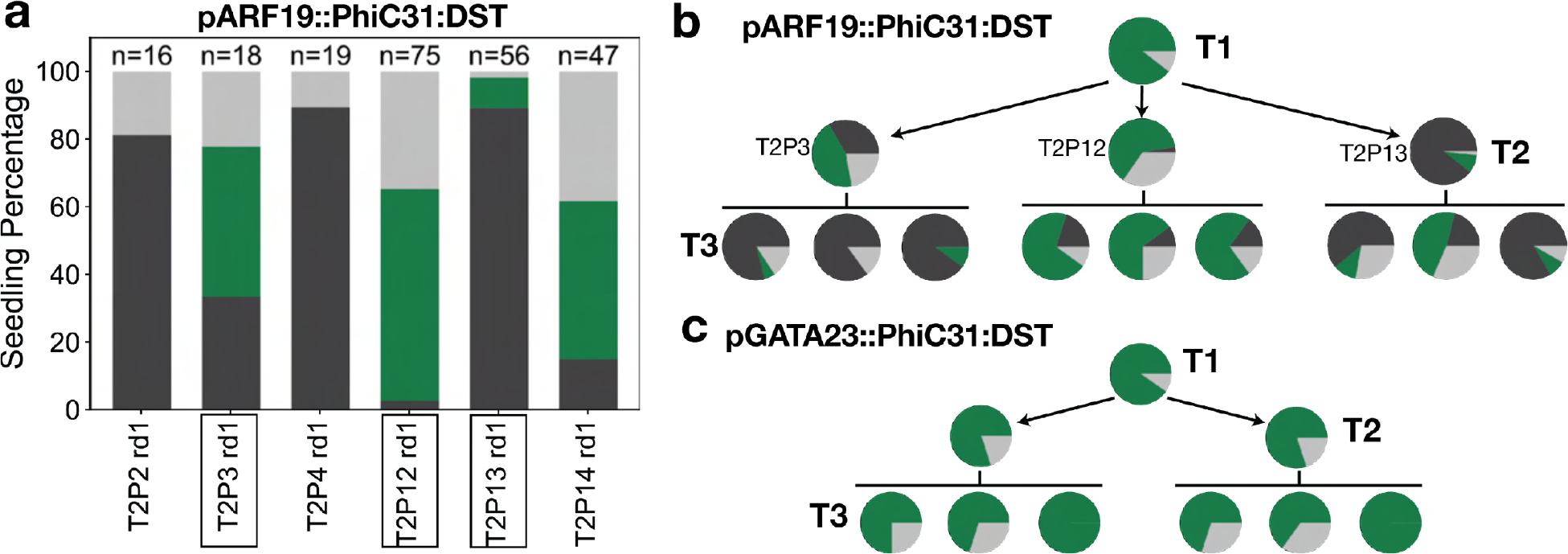
Additional data for pARF19::PhiC31:DST and pGATA23::PhiC31:DST (Figure 4). (a) Phenotyping of T2 seedlings from T1 lines with pARF19::PhiC31:DST in PhiC31 target line. The graph corresponds to the percentage of seedlings in each of the defined phenotype categories, such as no switch corresponding to no mScarlet expression in the root, switch in LR only: mScarlet expression only in the lateral root, switch not exclusive to LR: mScarlet expression in the main root. The number of seedlings characterized for each construct is mentioned at the top of the bar in the graph. The lines used in Figure 4 have their name boxed. (b) and (c) Phenotype of T1, T2, T3 plants from pARF19::PhiC31:DST (b) and pGATA23::PhiC31:DST (c) constructs in PhiC31 target line. The pie charts are another representation of the previous phenotype bar graph representing the percentage of seedlings in each of the defined phenotype categories. From each generation, three seedlings with the LR-only switch phenotype were kept to generate the next generation.

**Fig S8:**
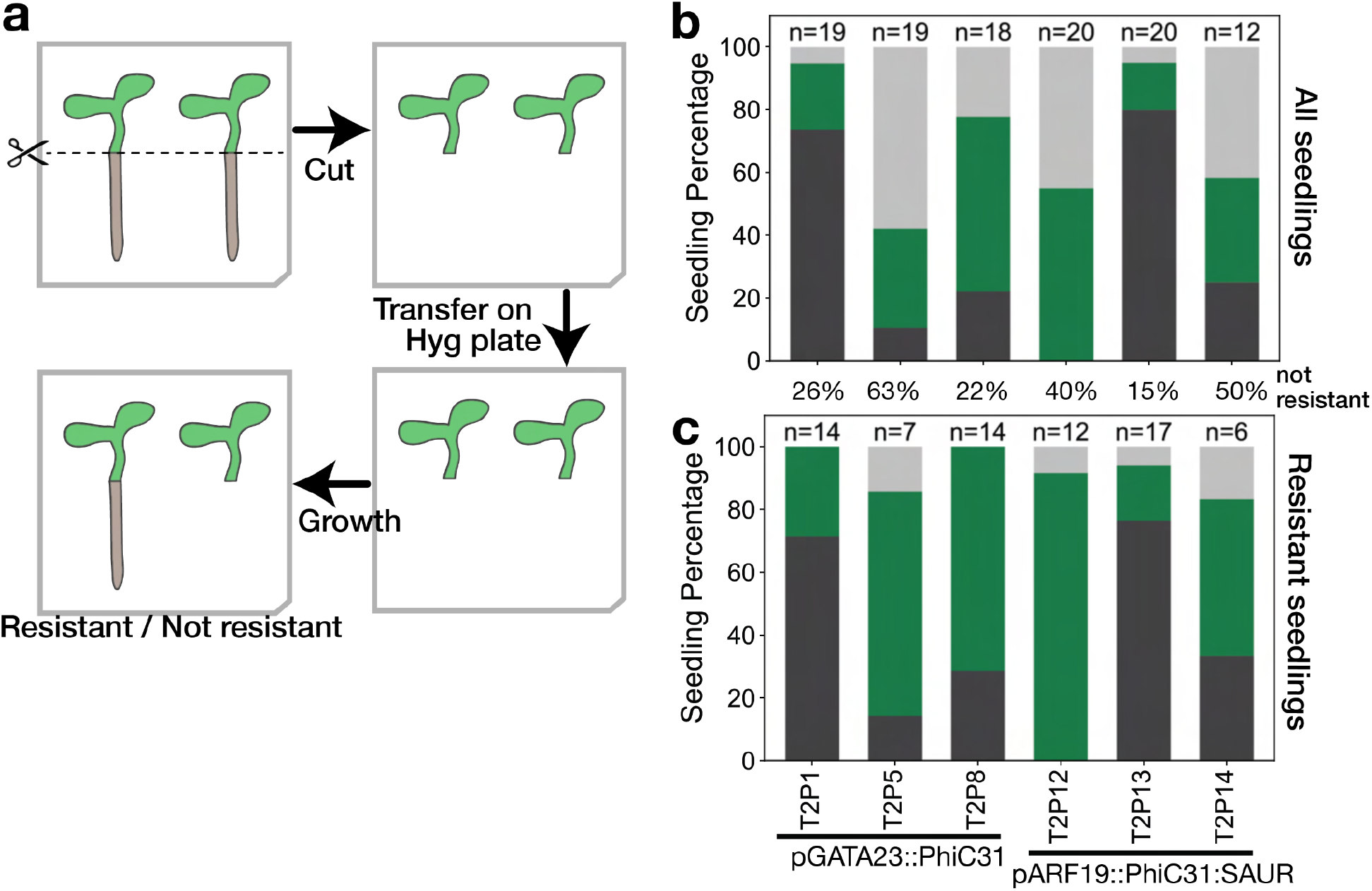
Post-phenotyping selection of seedlings (Figure 4). **(a)** Process to determine post-characterization if a seedling is resistant to hygromycin. Post microscope characterization, seedlings were cut between the hypocotyl and the root. The aerial tissue (hypocotyl and cotyledons) were transferred to a hygromycin plate and grown for seven days. Resistant seedlings grew new roots. (b) and (c) Phenotype of T2 seedlings for pGATA23::Phic31 and pARF19::PhiC31:SAUR constructs in PhiC31 target line. Labels of the lines are at the bottom, T2PX corresponds to the name of the specific line. (b) is the phenotype characterization of one round of T2 seedlings for those lines (number at the top of the bars). (c) is the same data from b with only Hyg resistant seedlings included. The percentage of seedlings which are not resistant is represented between both graphs. The graph corresponds to the percentage of seedlings in each of the defined phenotype categories: no switch (light gray); LR only (green); not exclusive (dark gray).

**Fig S9:**
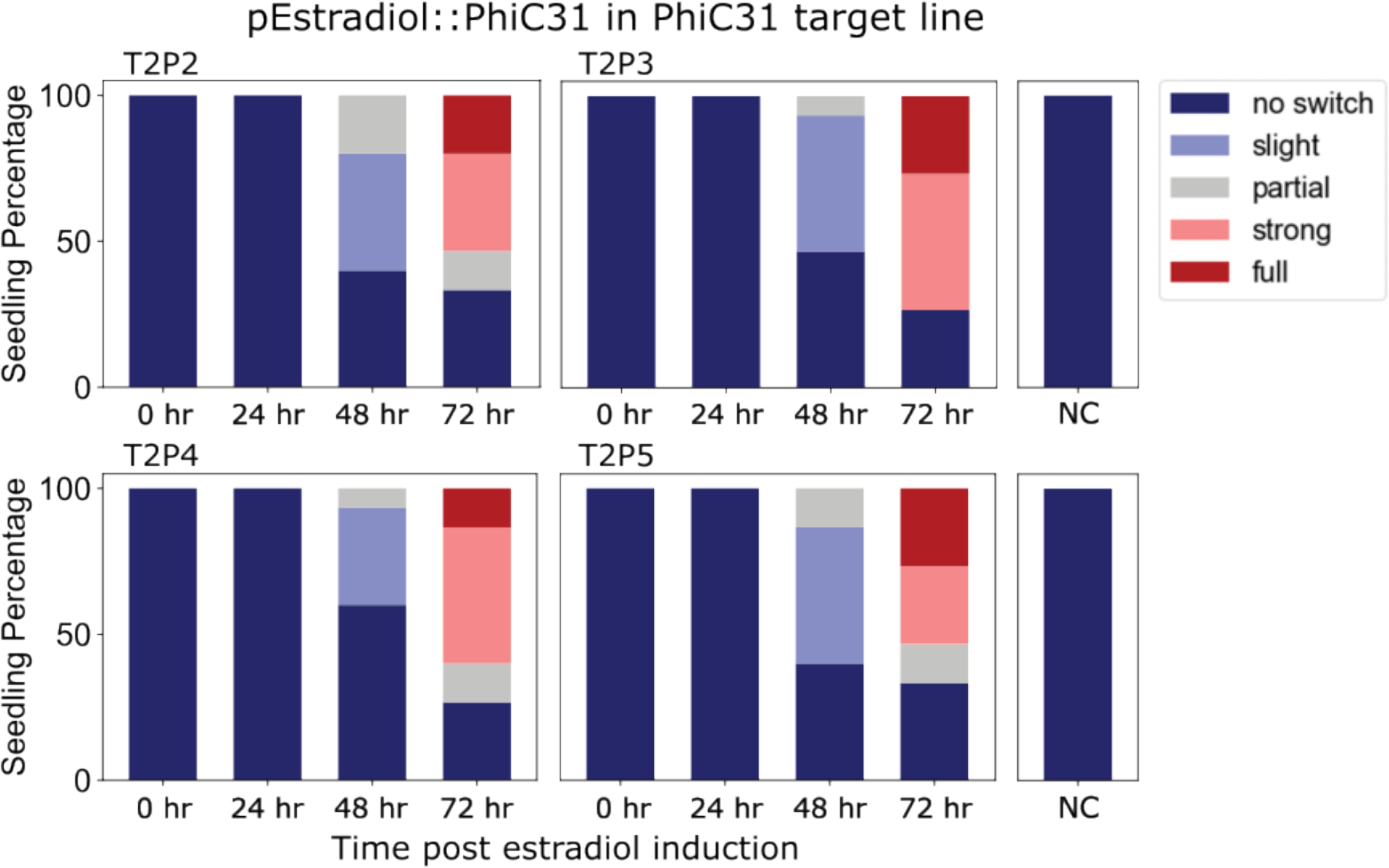
Estradiol induction of PhiC31 integrase in T2 seedlings. T2 seeds were collected from T1 plants which were not treated with estradiol. Four T2 lines were tested: T2P2, T2P3, T2P4, and T2P5 with 15 seedlings screened for each line. Seedlings were imaged and classified based on switching level as per Fig S5 at 0hr, 24hr, 48hr, and 72hr post-estradiol induction.

**Fig S10:**
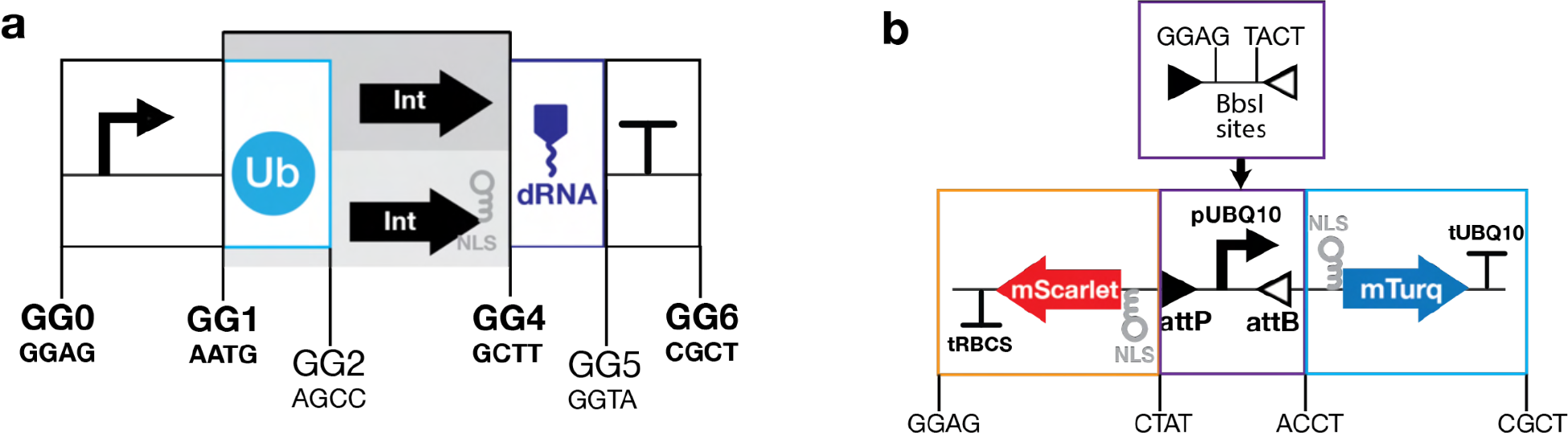
Cloning strategy based on golden gate assembly. (a) Cloning strategy for the integrase construct with the BsaI spacer between each part specified. (b) Cloning strategy for the integrase construct. The central part with the integrase sites and promoter is constructed by golden gate assembly with BbsI enzyme to add the promoter to the synthetic fragment with the integrase sites only.

## Supplementary Material and Methods: Python and ImageJ code

### a ImageJ macros to automatically analyse fluorescent microscope images

**Prior to use the micro:**

This macro allows you to take a .lif file and automatically process many multichannel images from the fluorescent microscope as a batch. It adjusts brightness/contrast, adds appropriate lookup tables, merges channels, and adds a scale bar for 10x magnification. Dependency: Save_all ImageJ plugin can be found https://imagejdocu.list.lu/plugin/utilities/save_all/start

Instructions for macro use

1. Import .lif file. In the pop-up check only the Autoscale box. Set “view stack with” as Hyperstack and set “Color mode” as colorized.
2. Hit Ok and select desired image set.
3. Go to Plugins>SaveAllImages and enter the directory where you want to save your processed images. For the “is this image a stack” option select No.
4. Go to Process>Batch>Macro. In the Input space enter where you saved your images in the last step. In the Output space put where you want your processed images to be saved. In the box, paste the macro from the macro .txt file in this repo.
5. Select process and your images will all be processed and in the specified output folder!

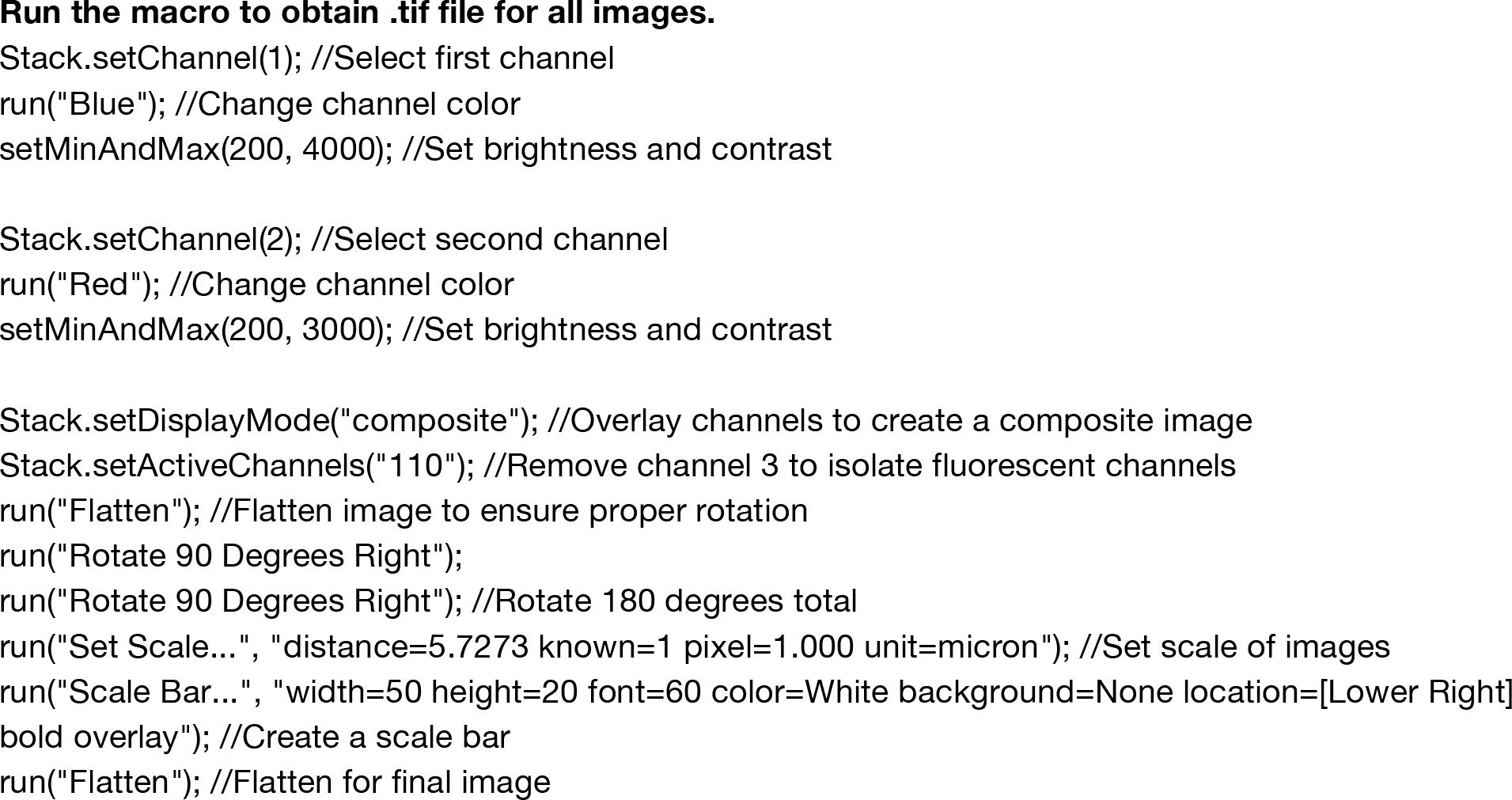

### b. Python script to generate bar graph for seedling characterization

This script allows you to create stacked bar plots showing the switching categories (either for tuning or lateral root switching data). It also computes statistical significance for the tuning dataset using analysis of variance (ANOVA) with a post hoc Tukey’s test.

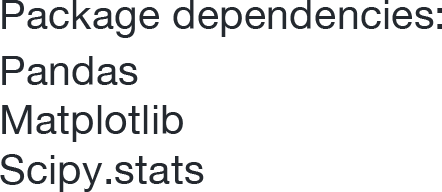
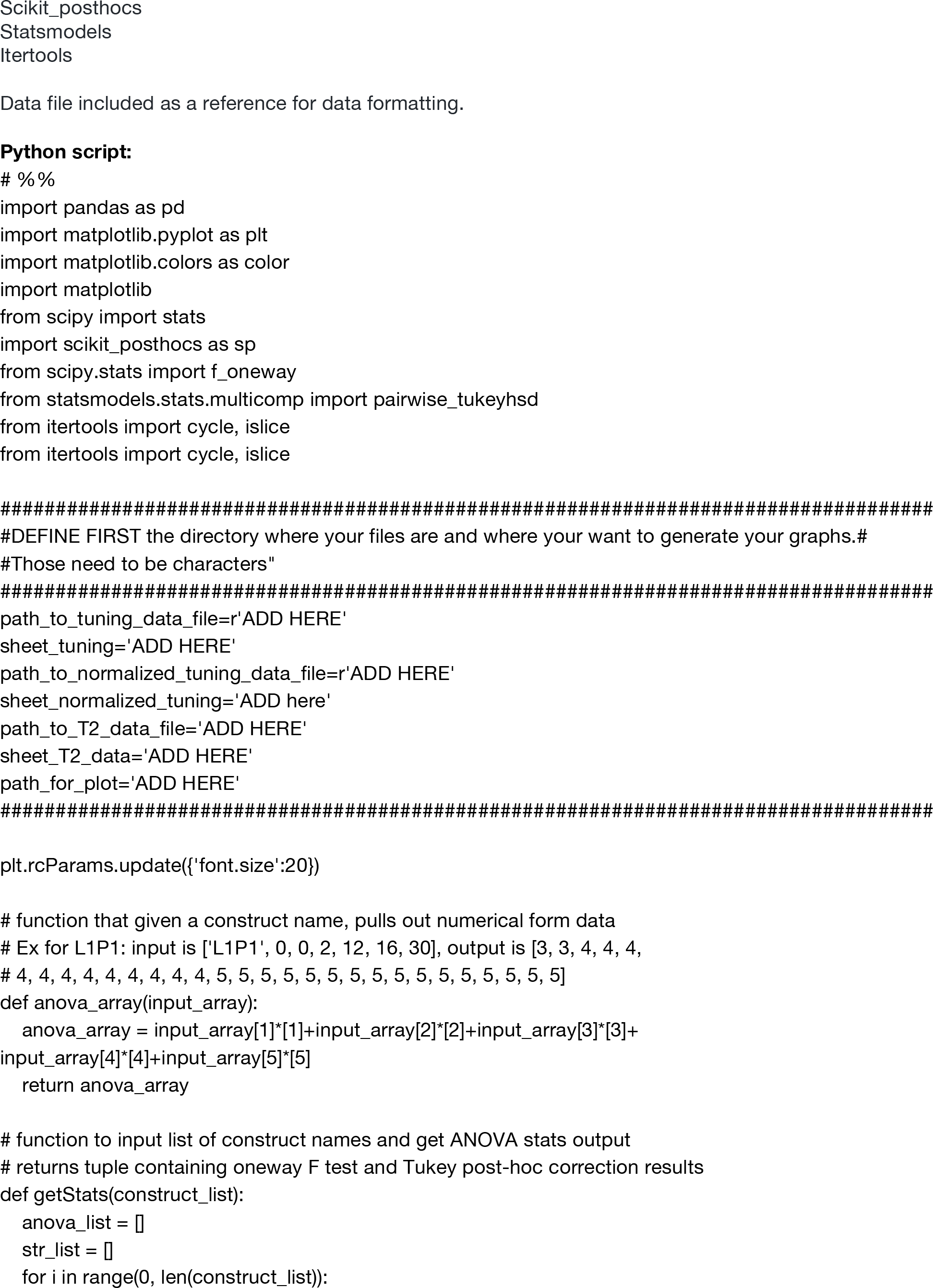
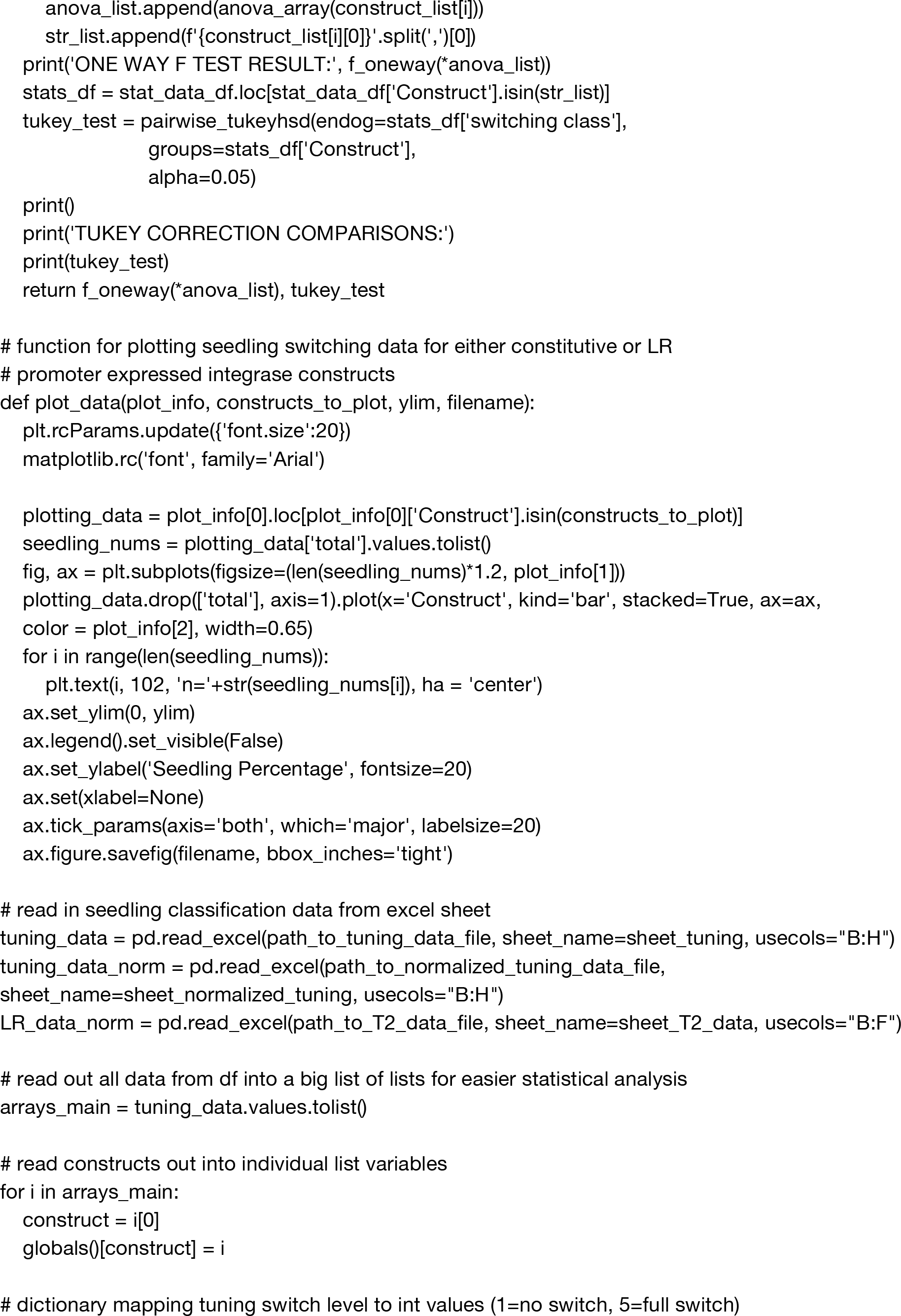
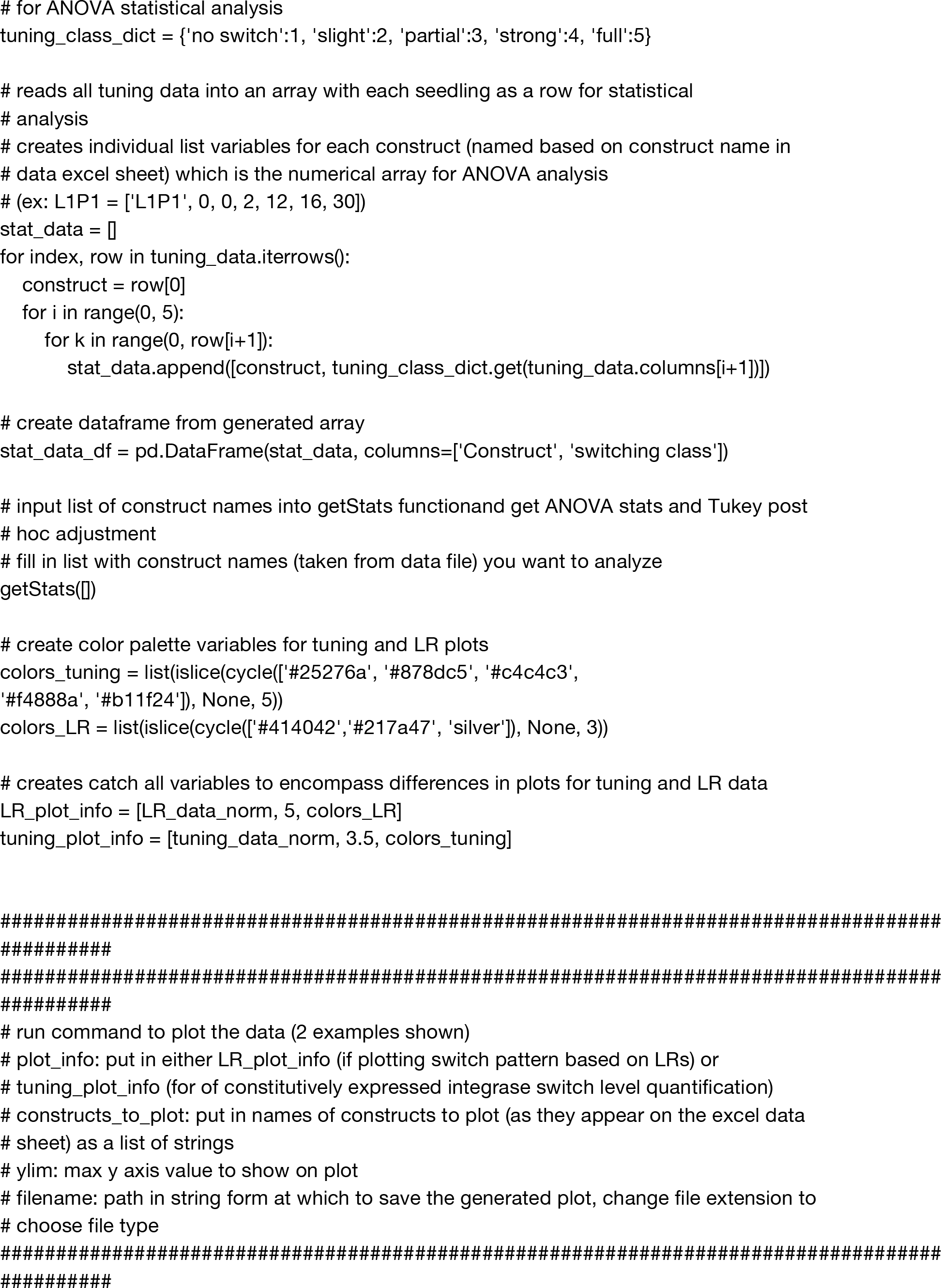
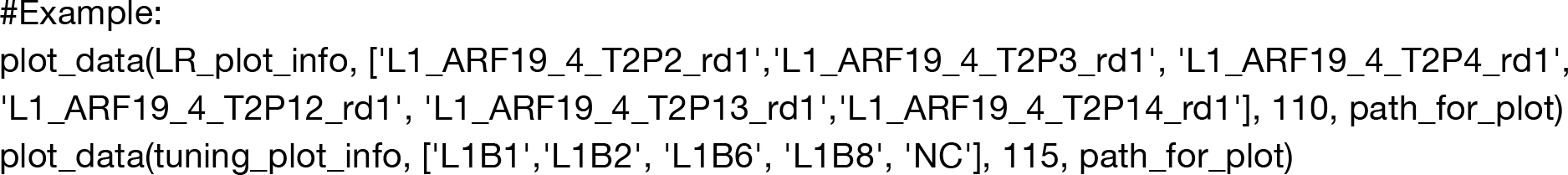

### c. Python script to generate plot for N. Benthamiana characterization

This script allows you to plot tobacco injection data (plate reader fluorescence measurements from injected leaf punches). It also computes statistical significance using ANOVA with post hoc Tukey’s test.

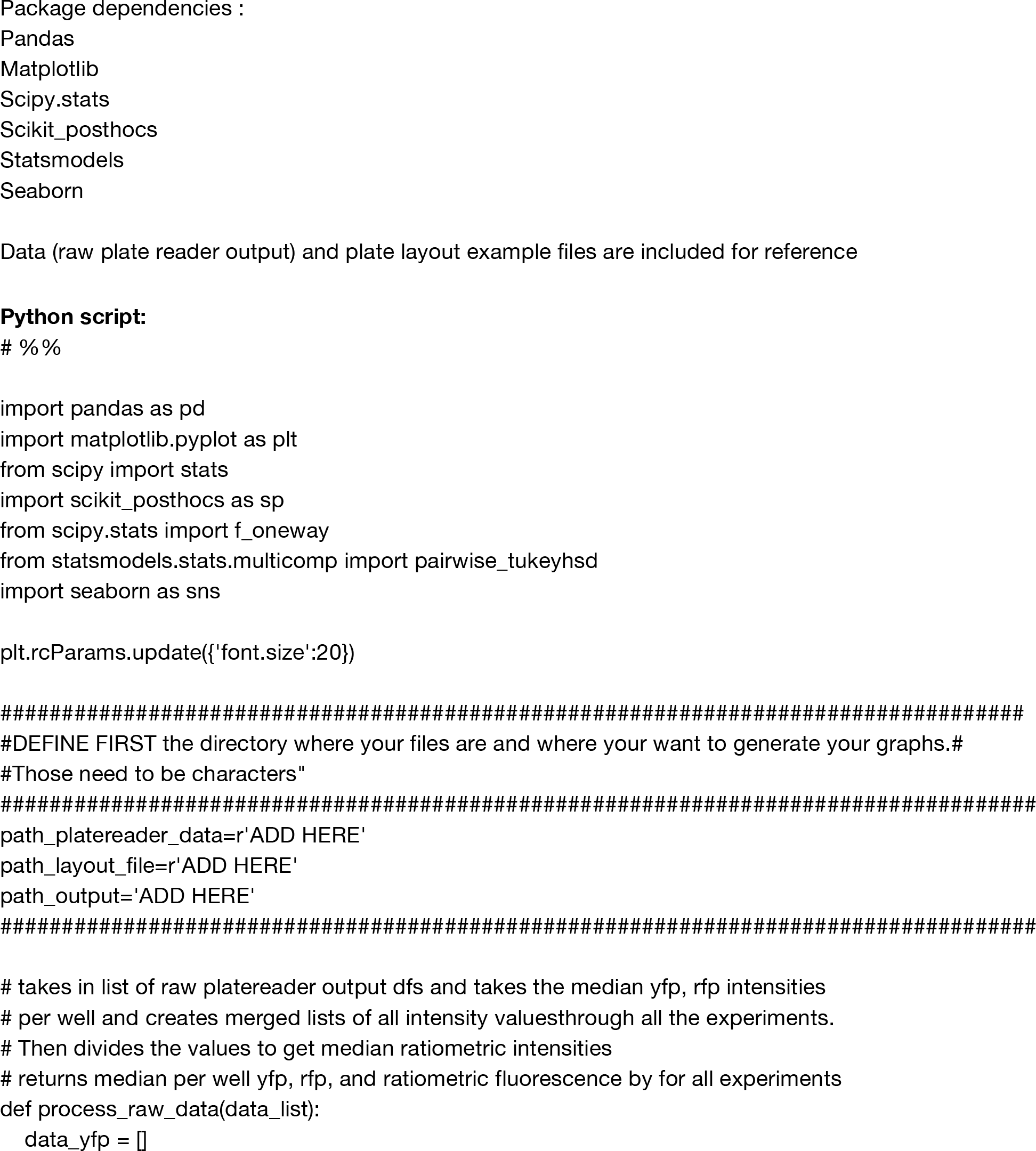
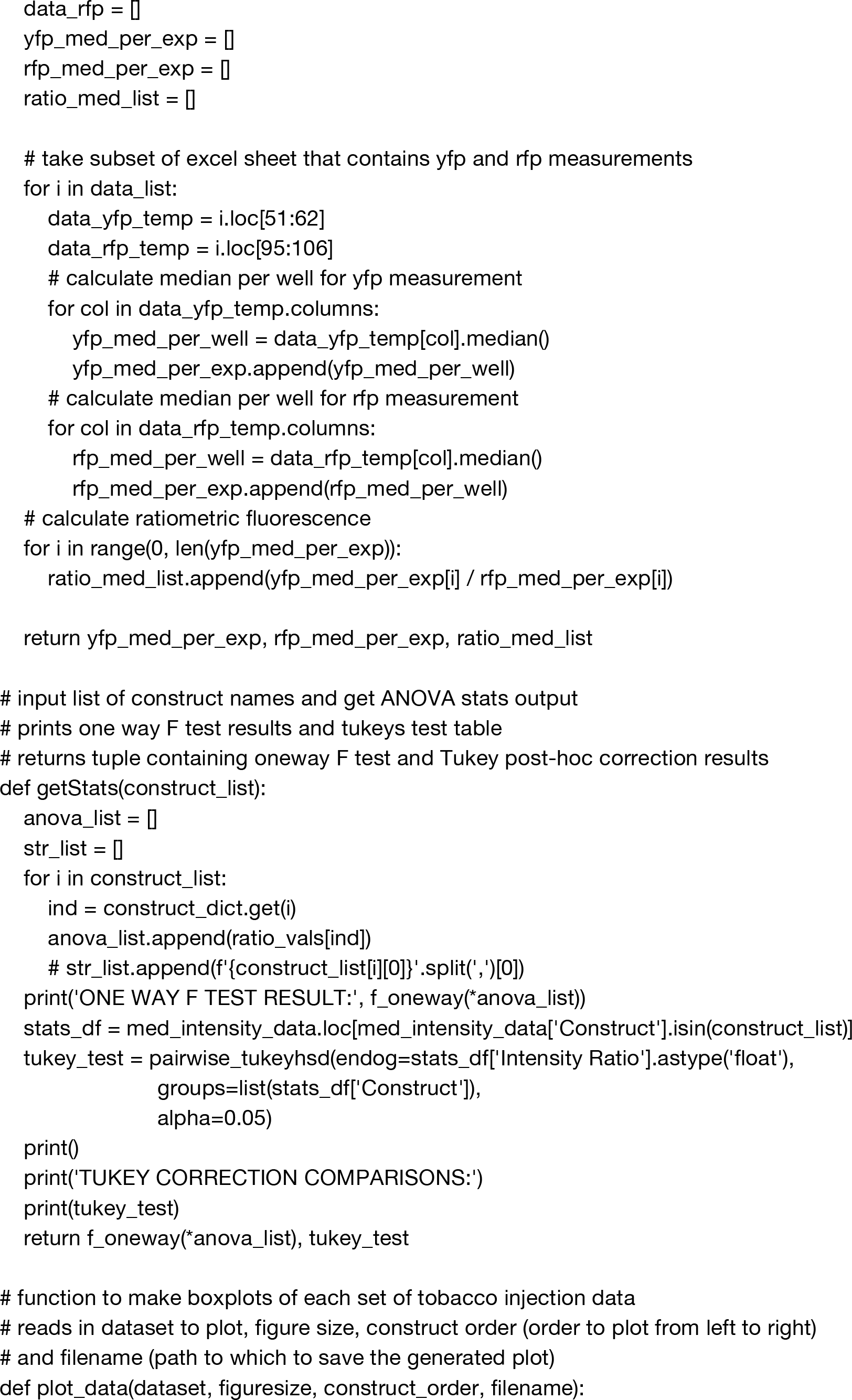
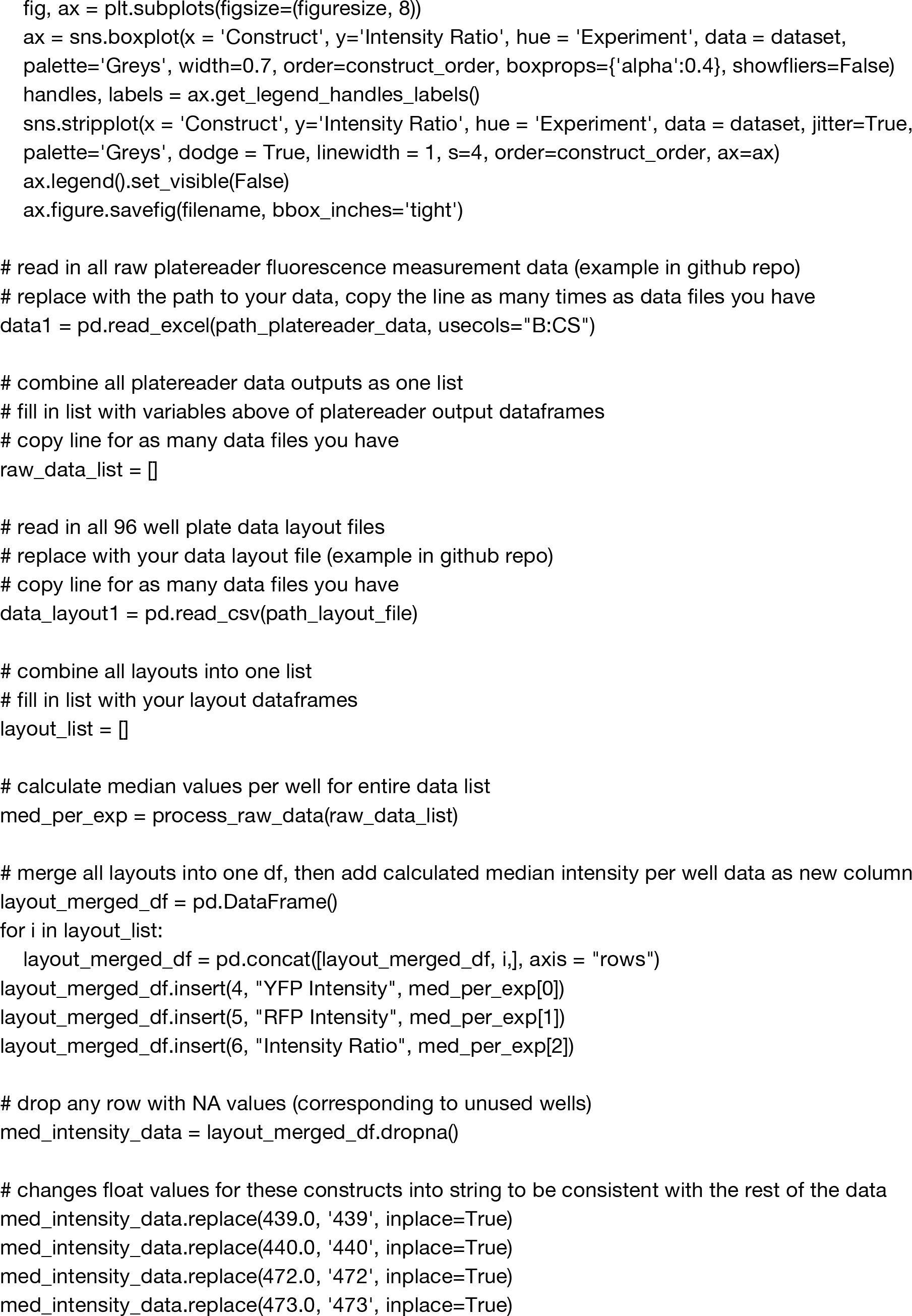
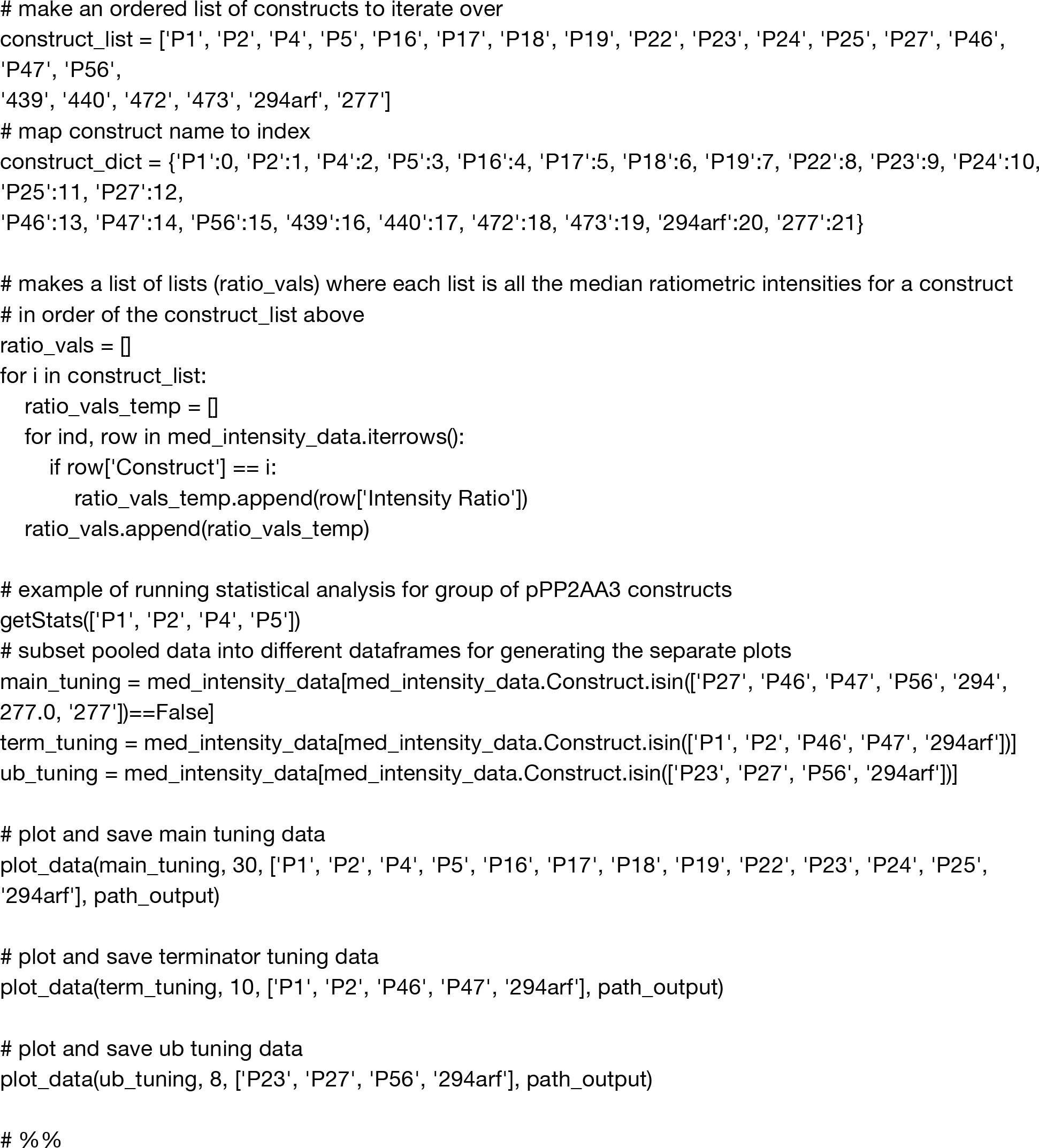

### Supplementary Dataset

**Dataset 1: Table of primers, constructs and plasmid sequences:** Available online (excel file).

**Dataset 2: Raw data for seedling characterization:** Available online (excel file).

**Dataset 3: Additional microscopy images.**

**Additional T1 images:** pDvp::PhiC31 and its tuning variants (+NLS and +DST) were each transformed into a plant line with an integrated PhiC31 target. Each T1 image is taken through the fluorescent microscope, overlaying the RFP and BFP channels to capture localization of integrase-mediated switching. Images are sorted into their phenotypic categories. “No switch” indicates a lack of integrase switching. “Switch in LR only” shows specificity in switching localized exclusively to the lateral root area. “Switch not exclusive to LR” shows a switch that occurs throughout the root, and is not only limited to the lateral root. Images in each category are placed alongside a color-coded bar corresponding to each category. Each image possesses an identification number placed in the top right corner of the image, linking it to a specific plant. Images from the same plant possess the same number but end with a unique letter.

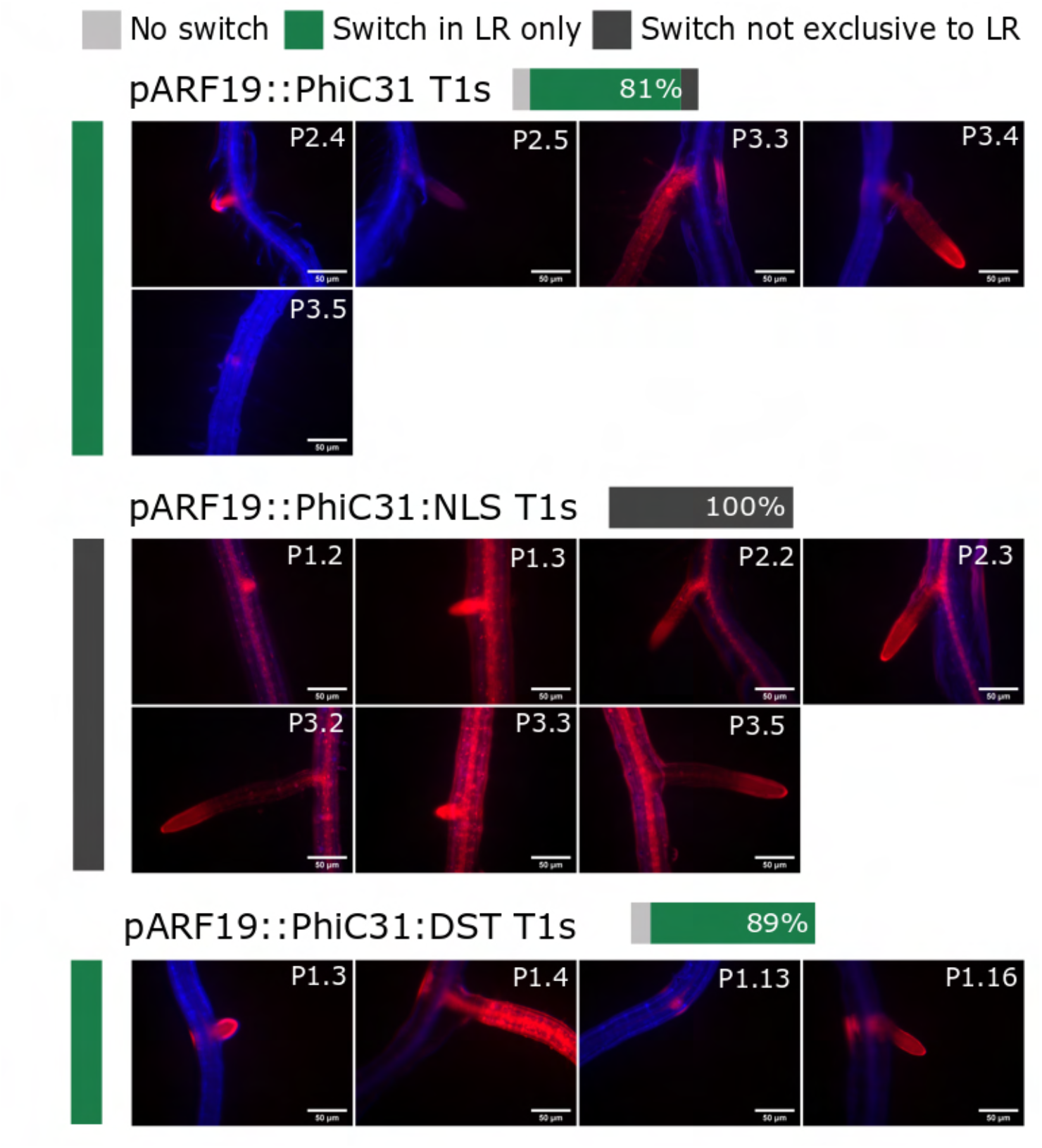
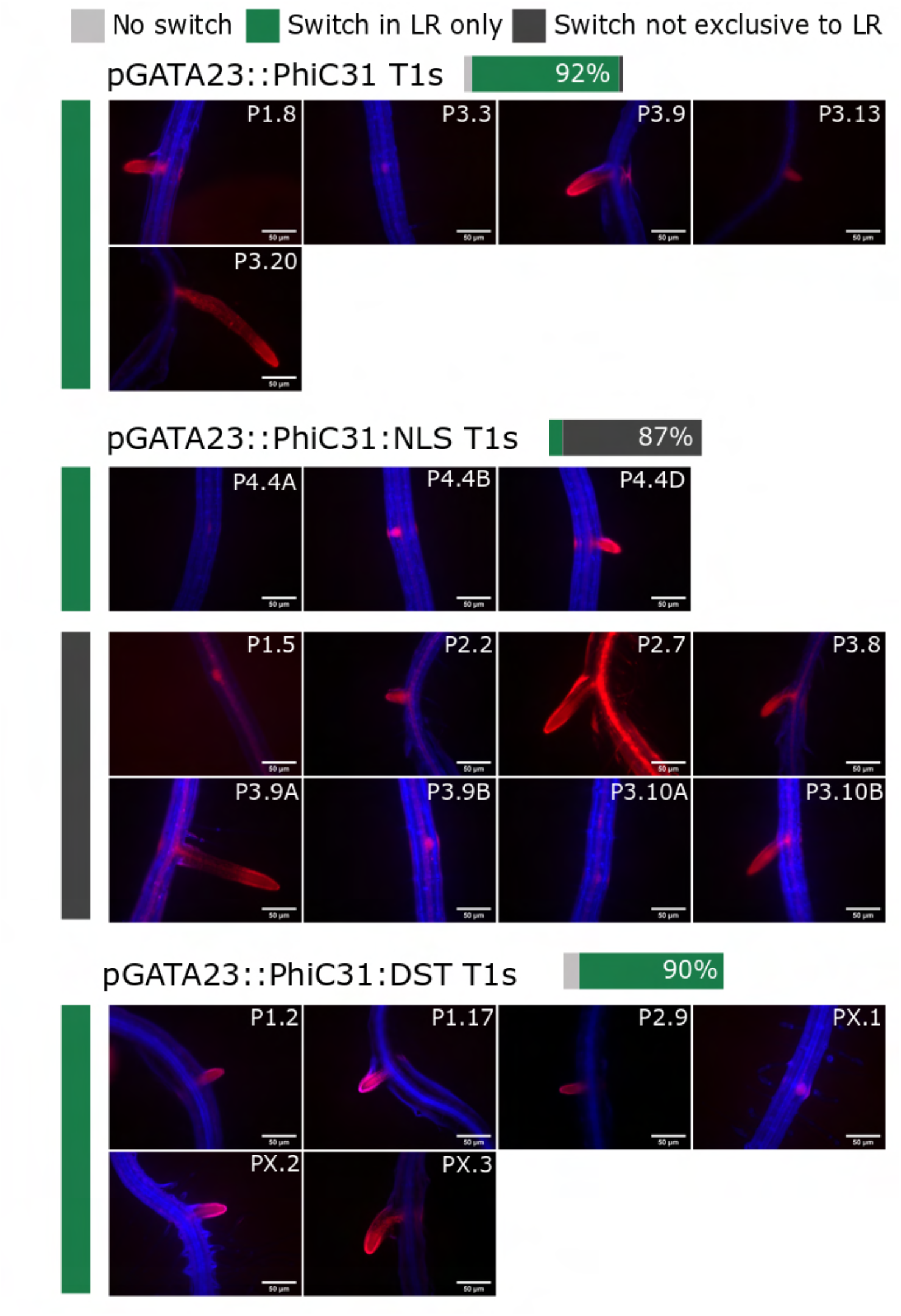
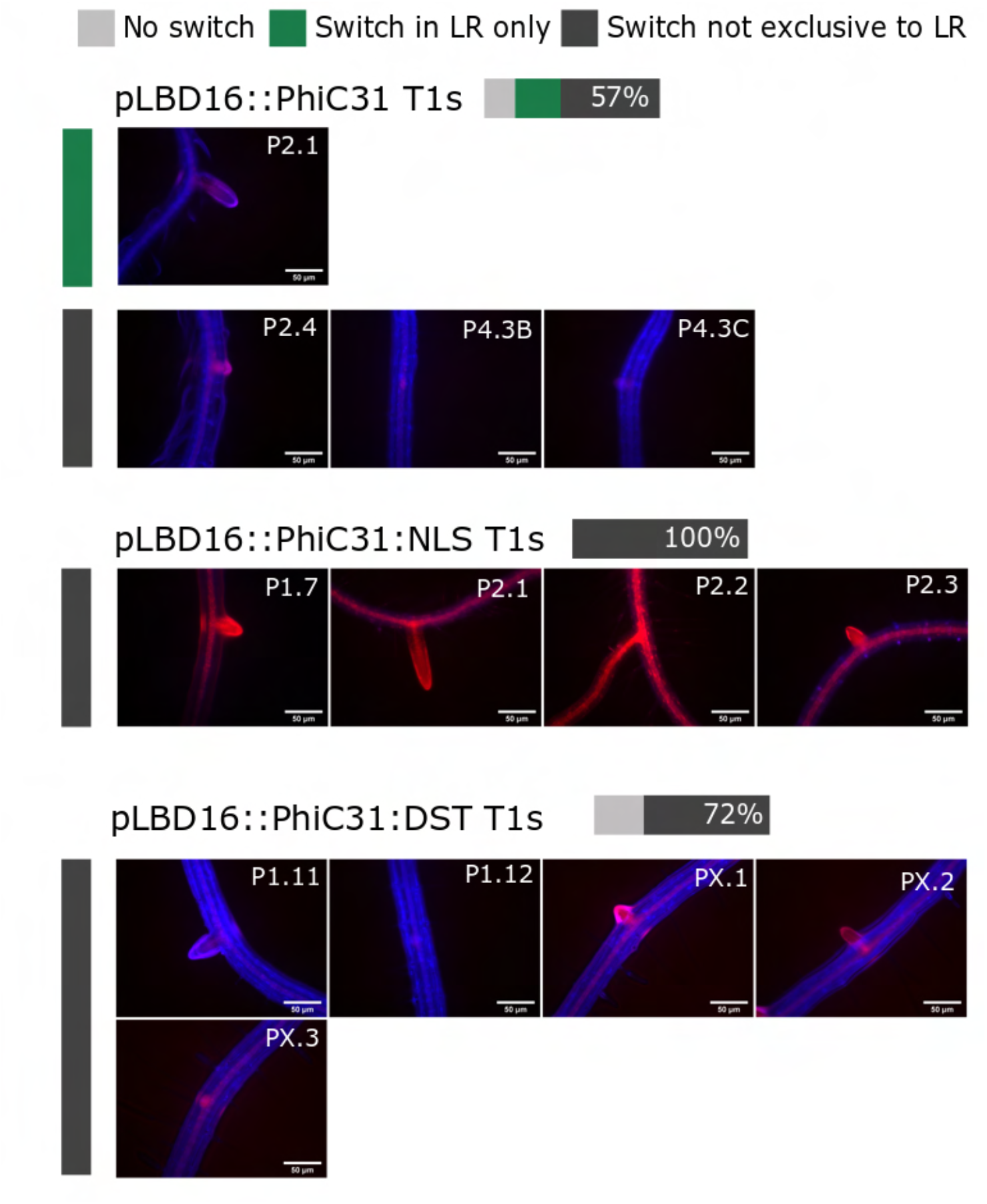

**Additional T2 images:** pDvp::PhiC31 and its tuning variants (+NLS and +DST) T2 seedlings were grown from the seeds of specific target T1 plants. Each T2 image is taken through the fluorescent microscope, overlaying the RFP and BFP channels to capture localization of integrase-mediated switching. Images are sorted into their phenotypic categories. “No switch” indicates a lack of integrase switching. “Switch in LR only” shows specificity in switching localized exclusively to the lateral root area. “Switch not exclusive to LR” shows a switch that occurs throughout the root, and is not only limited to the lateral root. Images in each category are placed alongside a color-coded bar corresponding to each category. Each image possesses an identification number placed in the top right corner of the image, linking it to a specific plant. Images from the same plant possess the same number but end with a unique letter.

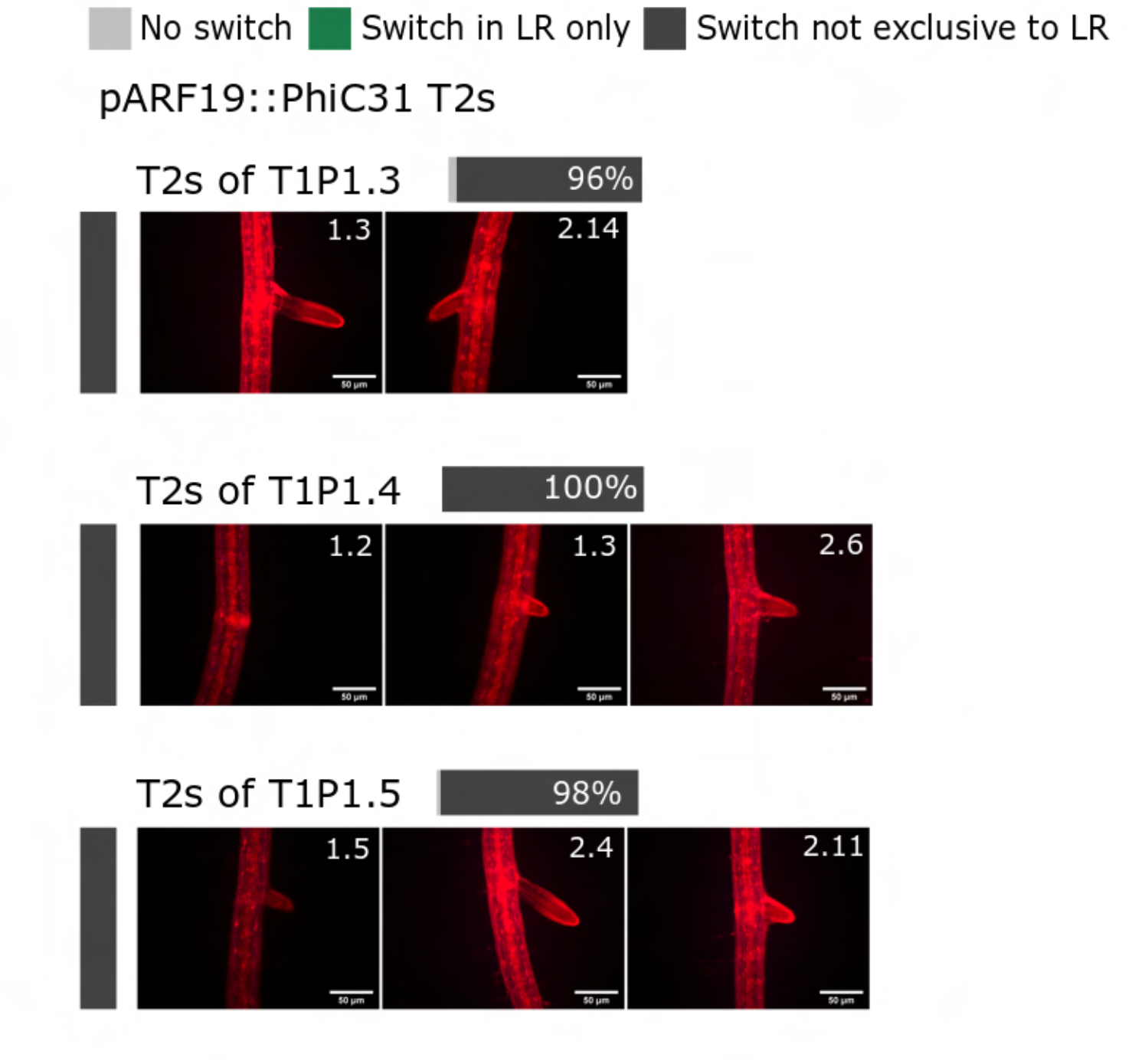
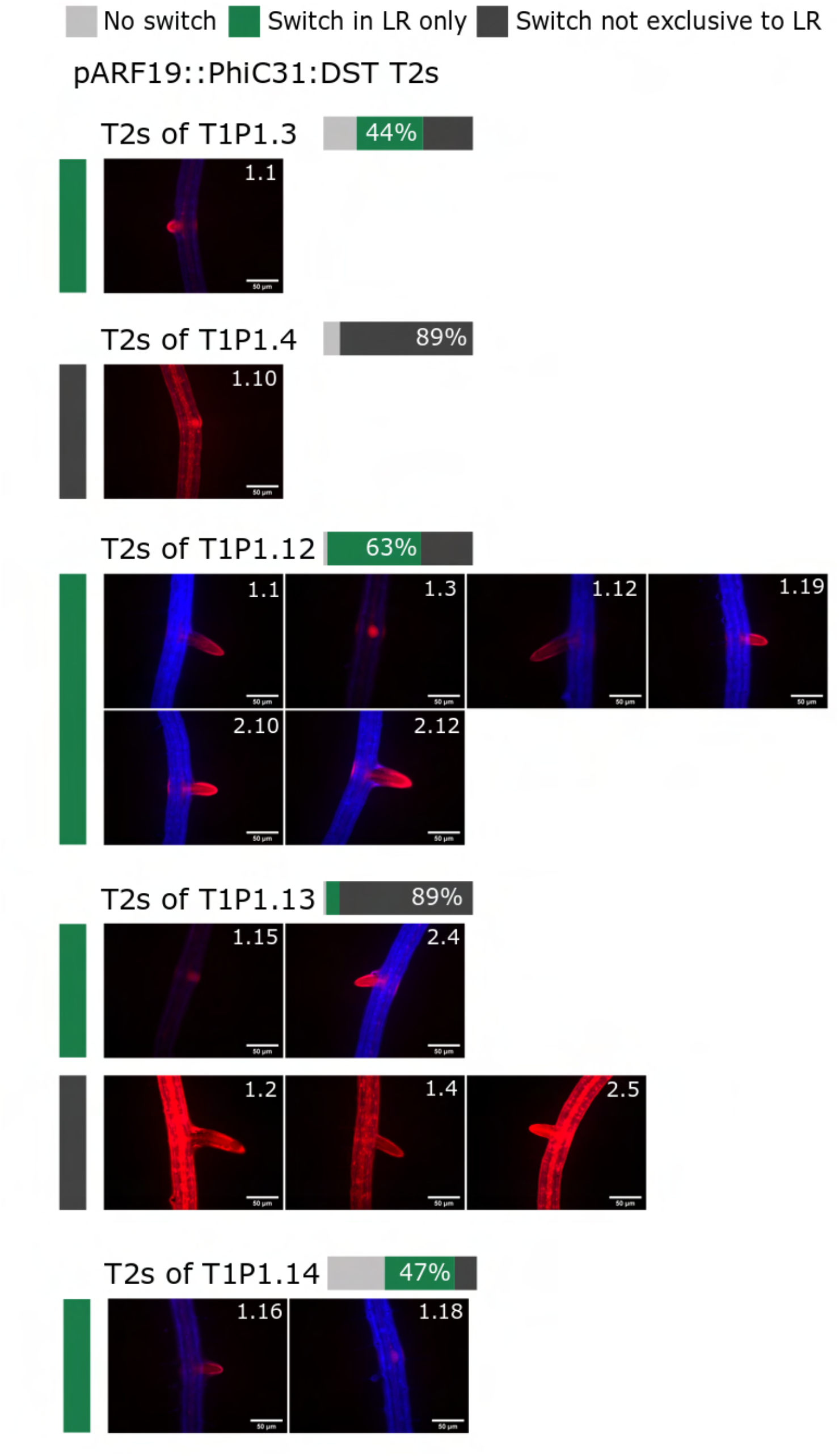
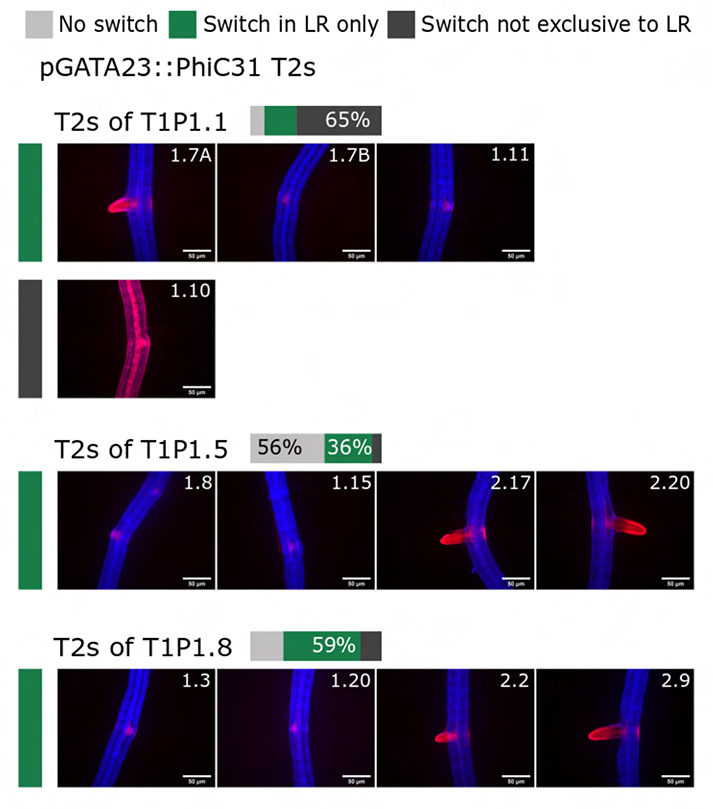
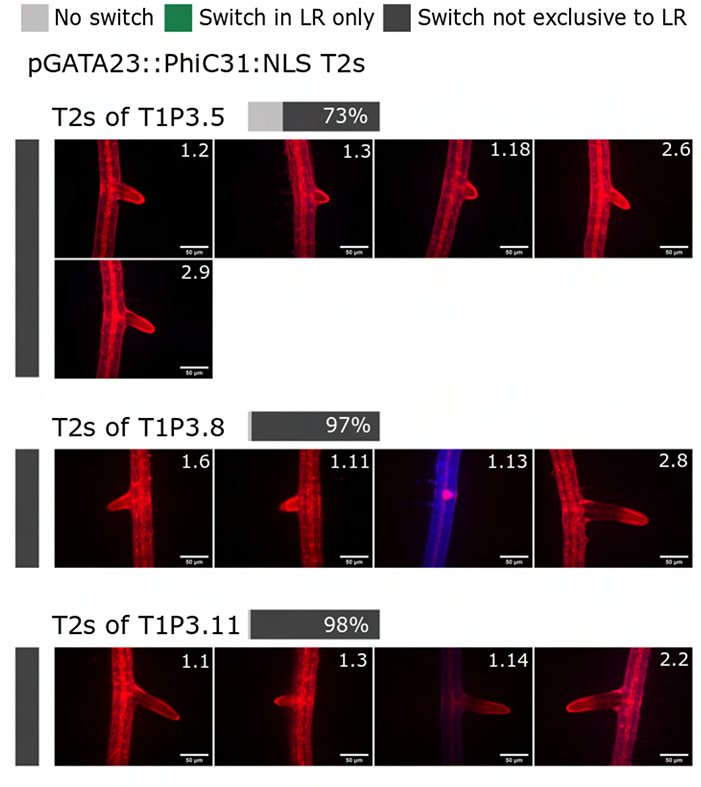
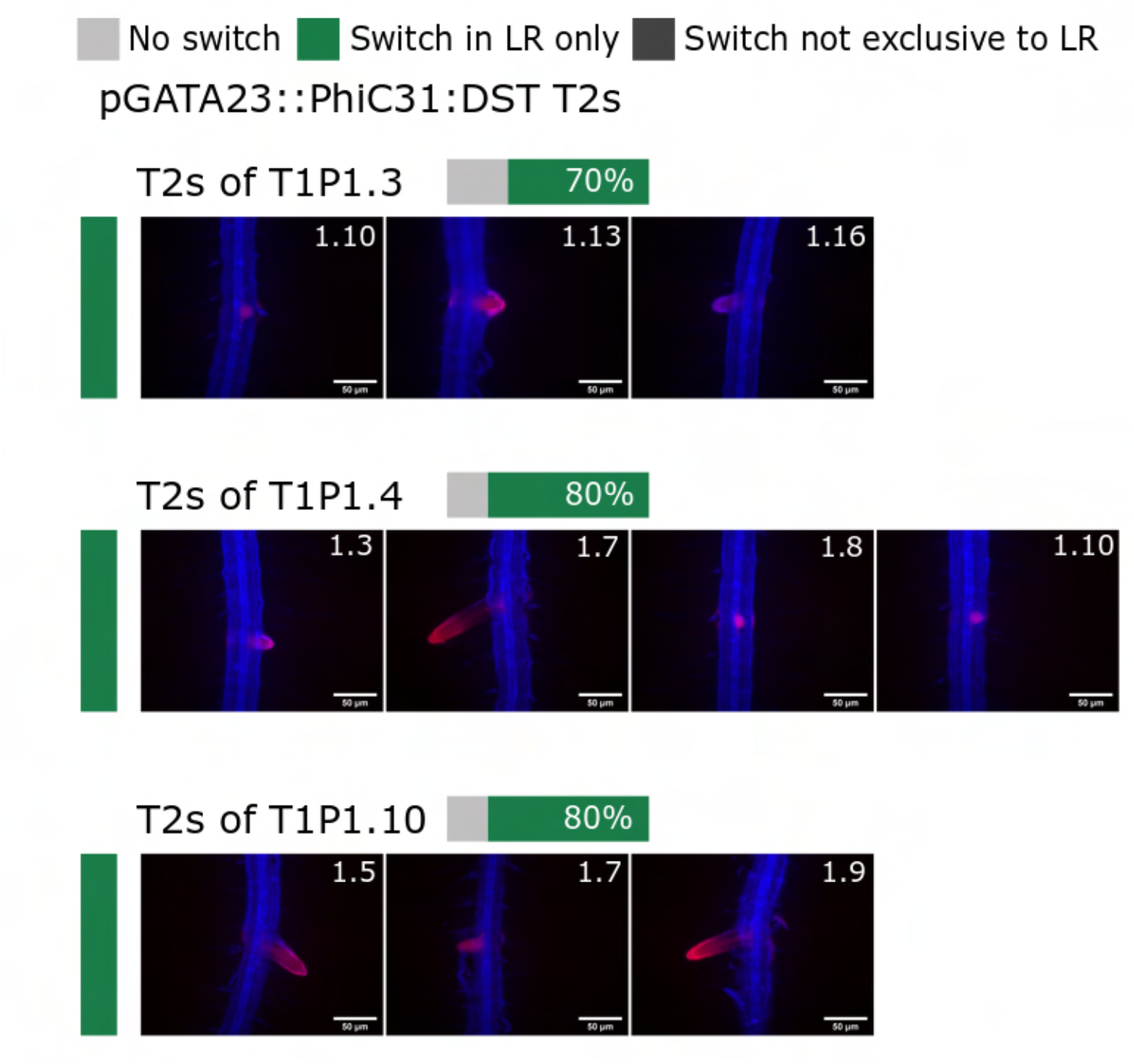
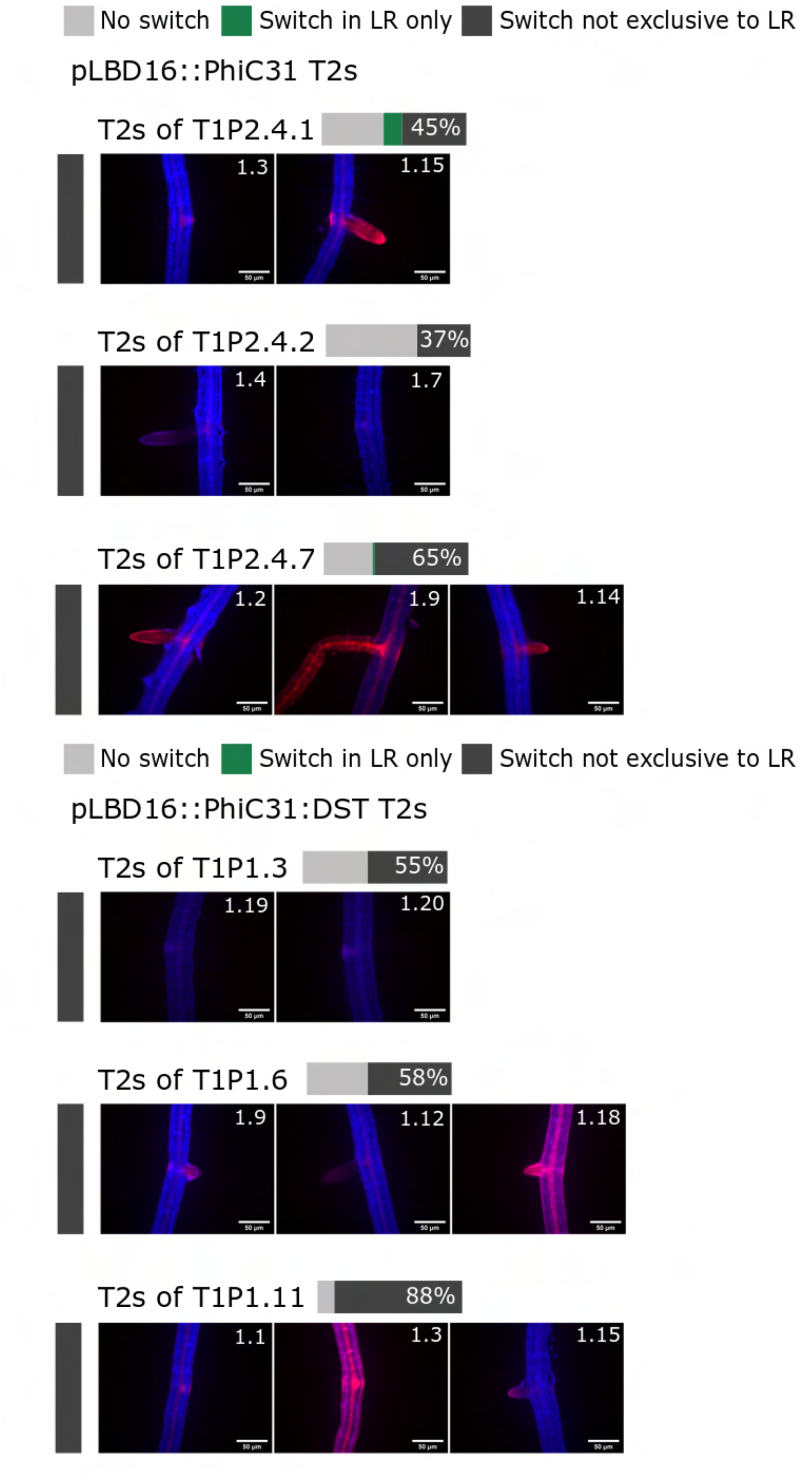

### Dataset4: Additional confocal microscopy images

**Confocal imaging of transcriptional reporters and integrase-based recorder in early-stage lateral roots**, corresponding to Fig 5 (more details in Fig 5 legend for plant lines). The top panel corresponds to an overlay of brightfield and red fluorescence channels from a single frame, the middle panel corresponds to the red fluorescence channel alone, and the bottom panel corresponds to the maximum projection of the Z-stack of seedlings in the red fluorescence channel.

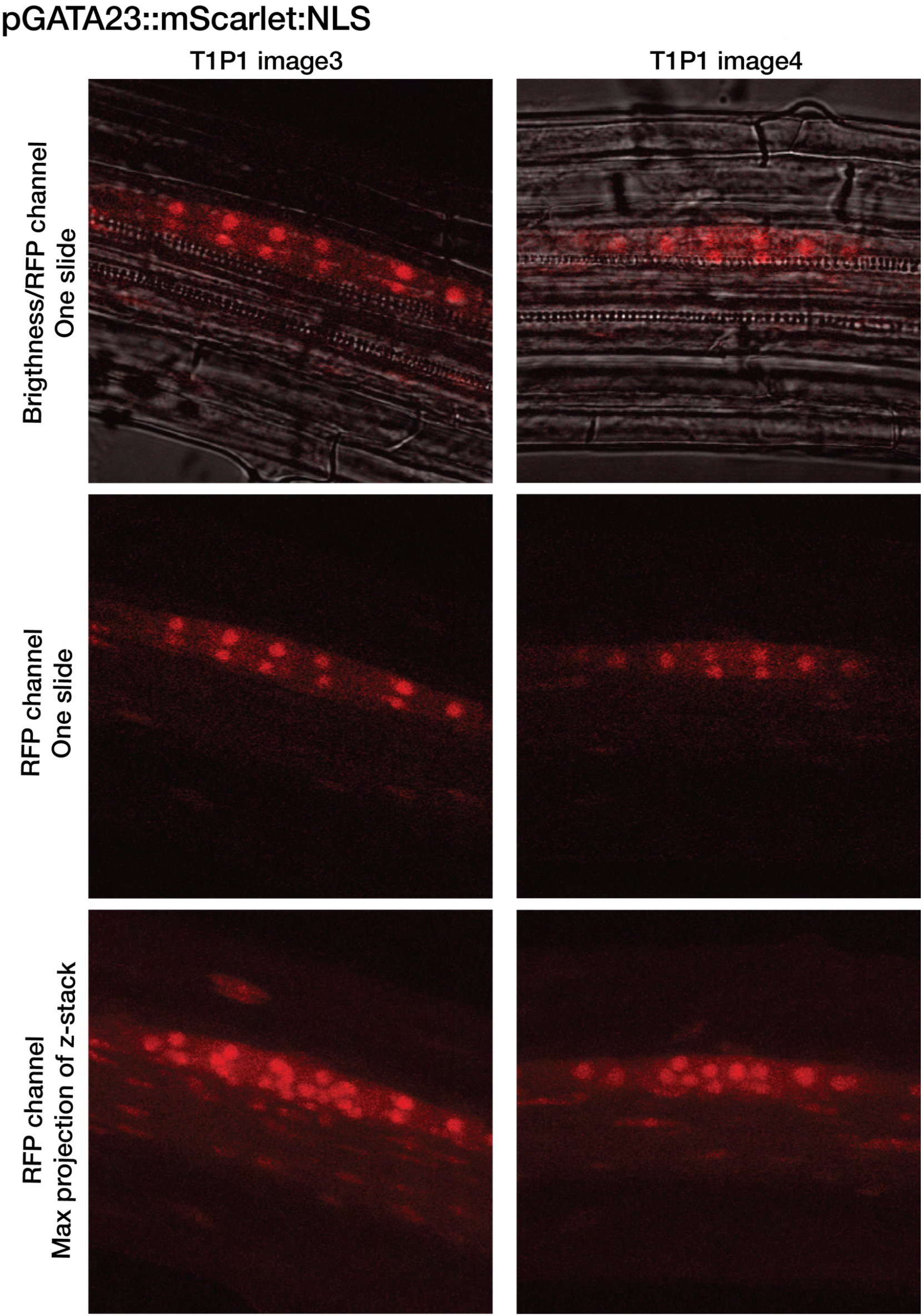
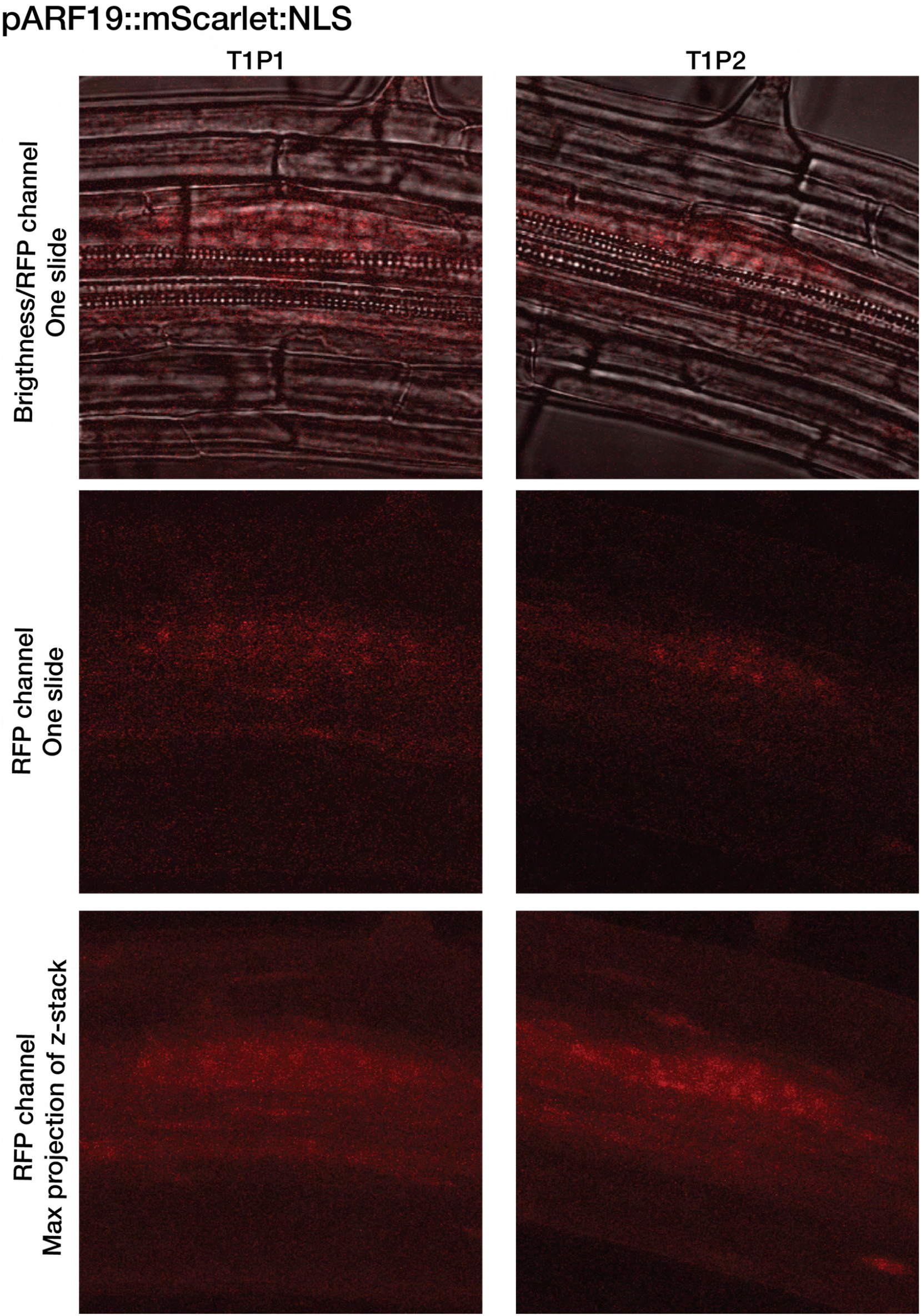
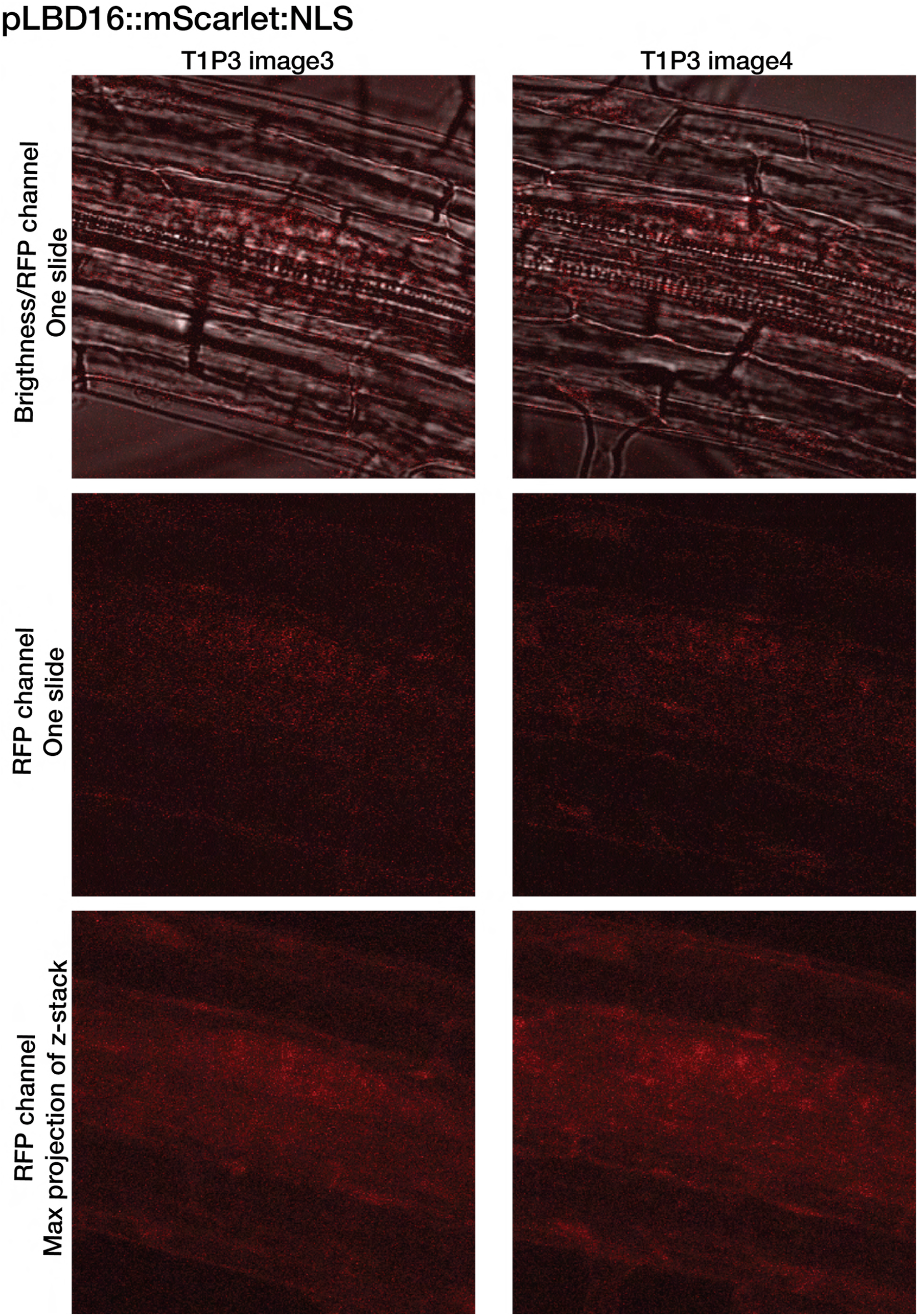
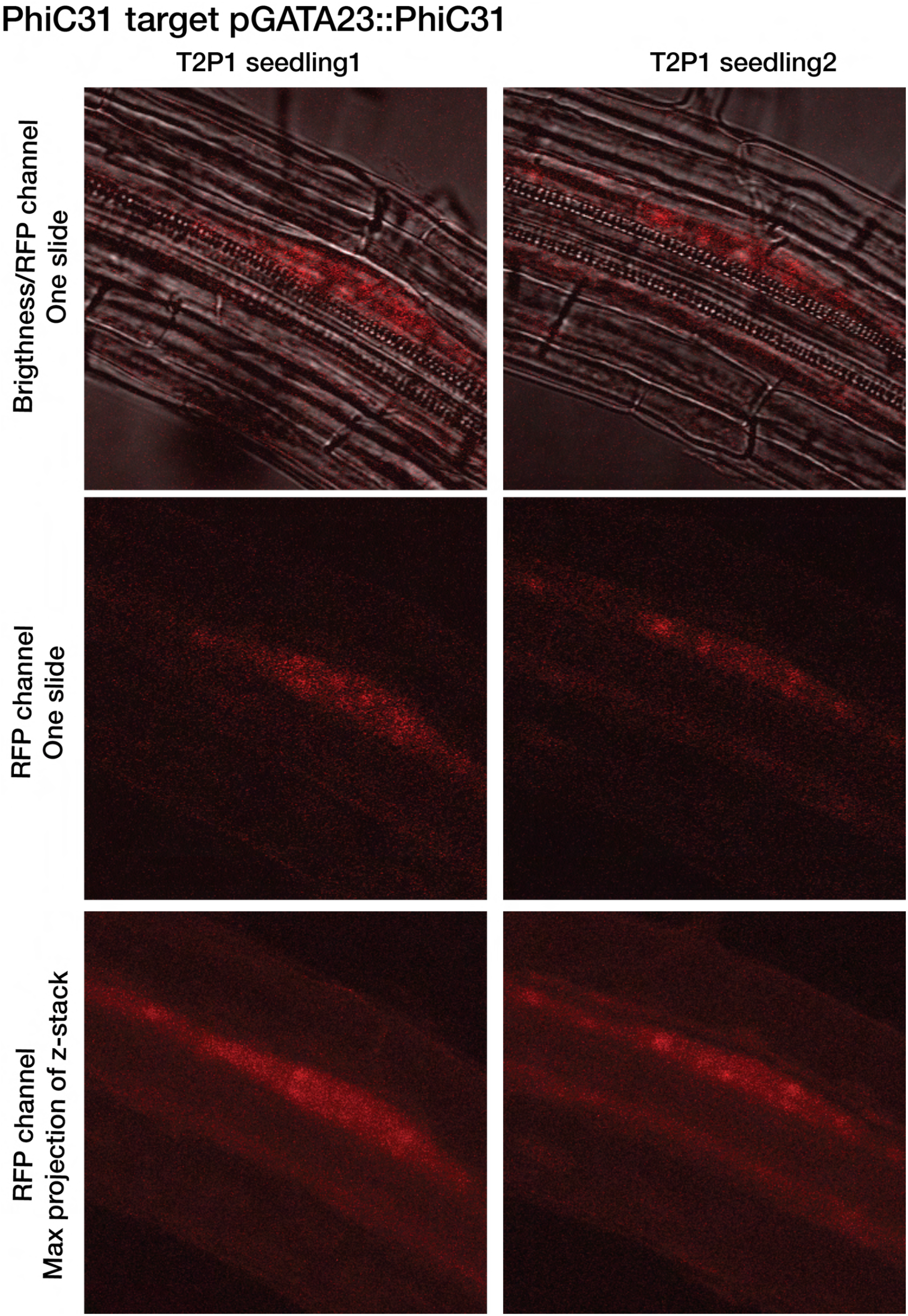
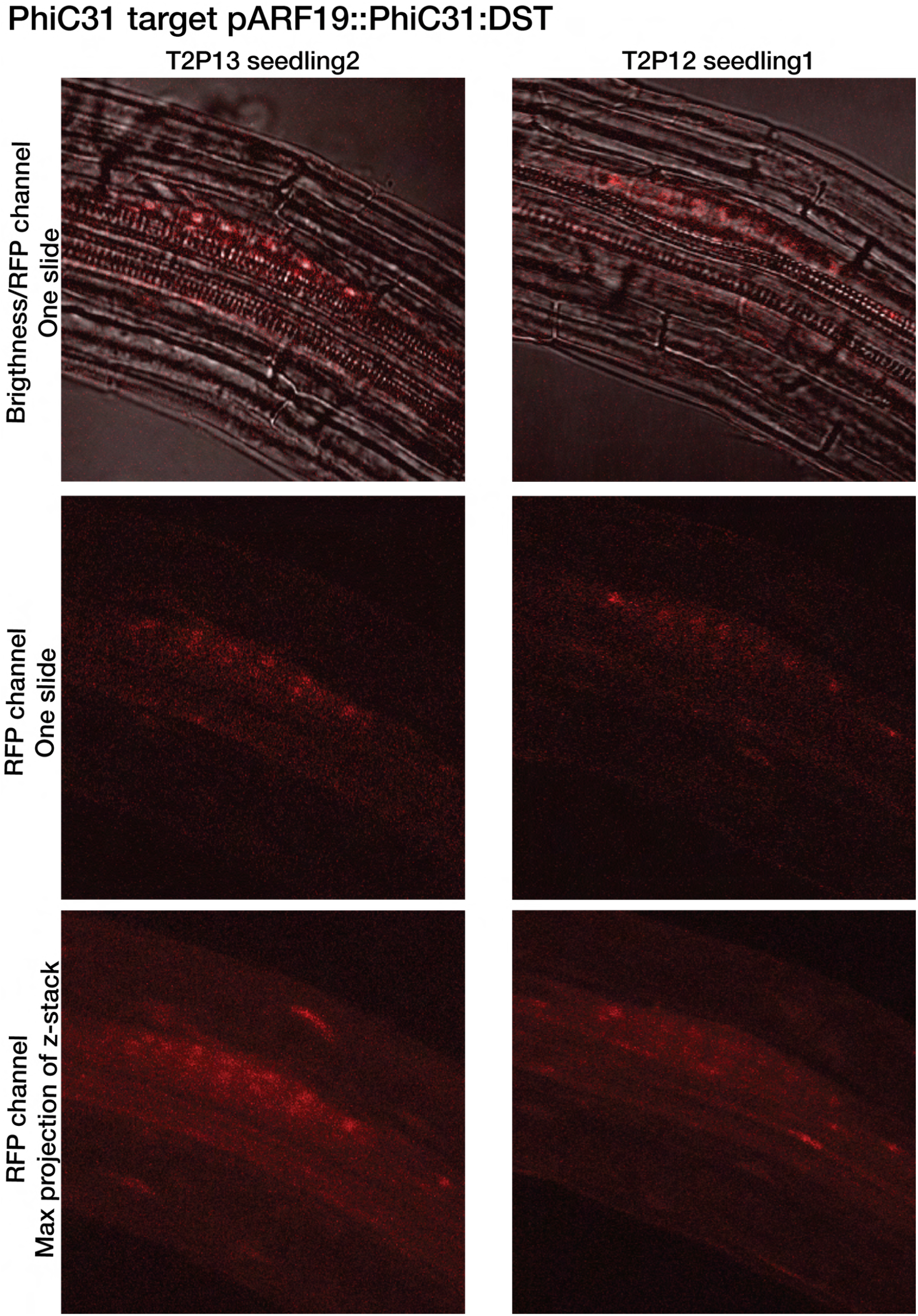
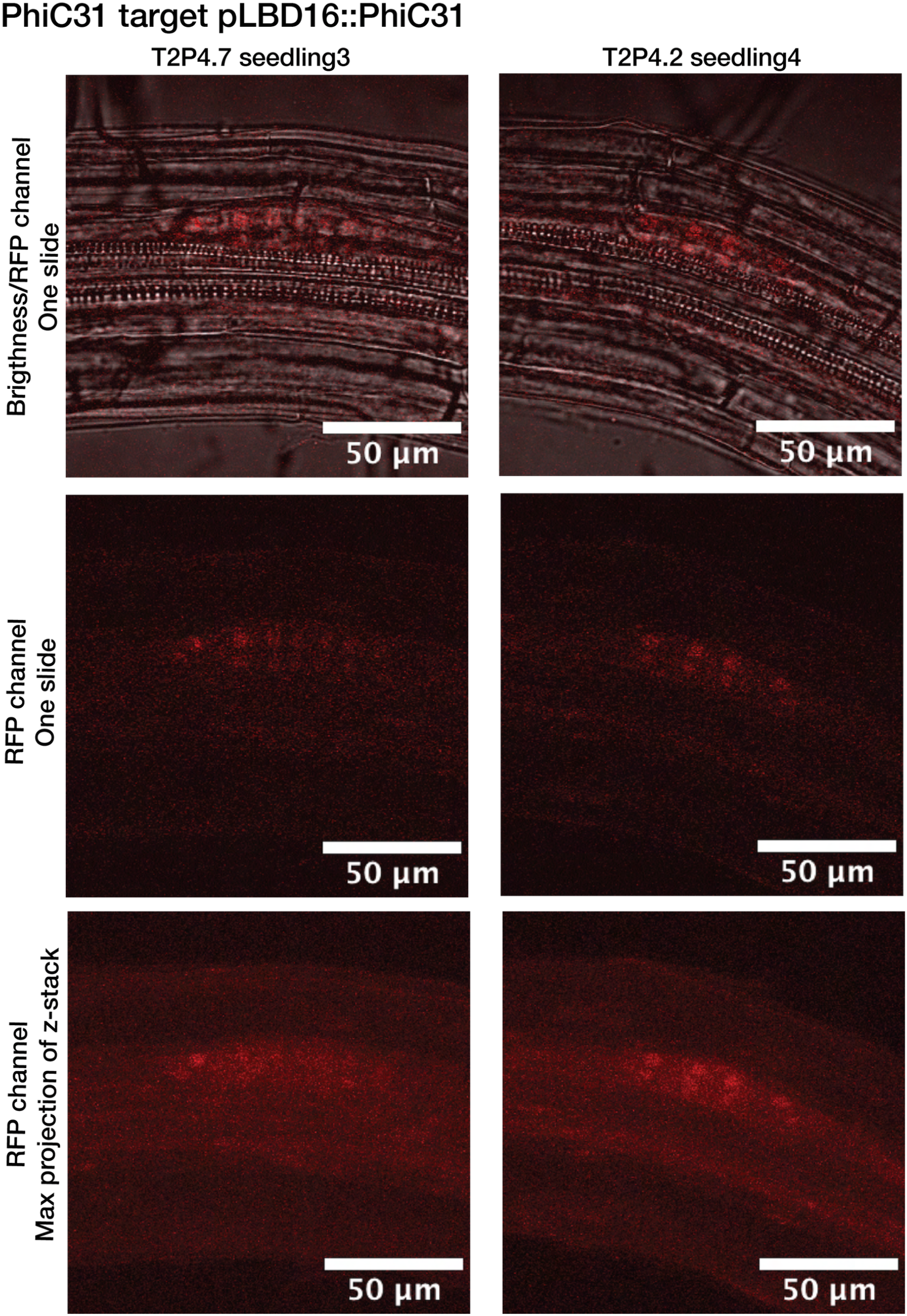

